# Absence of *Staphylococcus aureus* in wild populations of fish supports a spillover hypothesis

**DOI:** 10.1101/2022.10.18.512561

**Authors:** Marta Matuszewska, Alicja Dabrowska, Gemma G. R. Murray, Steve M. Kett, Andy J. A. Vick, Sofie C. Banister, Leonardo Pantoja Munoz, Peter Cunningham, John J. Welch, Mark A. Holmes, Lucy A. Weinert

## Abstract

*Staphylococcus aureus* is a human commensal and opportunistic pathogen that can also colonise and cause disease in other animal species. In humans and livestock, where *S. aureus* is most studied, there is evidence that strains have different host specialisms. Recent studies have found *S. aureus* in many wild animals, including fish, whose physiologies and ecologies are very different to humans. However, it remains unclear whether *S. aureus* is adapted to and persisting within these species, or if its presence is due to repeated spillover from a source population. Distinguishing between these two scenarios is important for both public health and conservation. In this study we looked for evidence to support the hypothesis that the presence of *S. aureus* in fish is the result of spillover, through testing for the presence of *S. aureus* in fish that are isolated from likely source populations. We sampled 123 brown trout and their environment from 16 sites in the Scottish Highlands. All these sites are remote and have very low populations density of wild animal species known to carry *S. aureus*, but were selected to represent variable levels of exposure to humans, avian and livestock species. While our sampling methods readily detected *S. aureus* from the external and internal organs of a farmed fish, we did not detect *S. aureus* in any wild trout or their environment from any of the 16 sites. We sequenced 12 *S. aureus* isolates from the farmed fish. While they were all from clonal-complex 45, the genomic diversity was high enough to indicate repeated acquisition from a source population. In addition, the presence of a φSa3 prophage containing a human immune evasion cluster indicates a recent history of these isolates within human populations. Taken together, our results support the presence of *S. aureus* in fish being due to spillover from other host populations, rather than the adaptation of *S. aureus* to aquaculture or fish populations. Given predictions that fish consumption will increase, more whole genome sequencing of *S. aureus* in aquaculture is needed to understand the presence of *S. aureus* in these environments and to mitigate the risk to fish and human health.

## Introduction

*Staphylococcus aureus* is both a common commensal bacterium of the human nasopharynx and skin, and an opportunistic pathogen (1). *S. aureus* also colonizes and causes infections in companion animals (2,3) and livestock (4,5) with the latter resulting in significant morbidity and economic loss (6). Recent studies have shown that major human and livestock strains, which are distinguished by a profile of house-keeping genes and named clonal complexes (CC), are also found in many wild animal species (7-12). However, little is known about whether this is a consequence of recent spillover from human and/or livestock populations (i.e., limited transmission of a strain in a novel host group with no evidence of adaptation), or if they persist independently within wild animal species. These two scenarios have different implications for both public health and conservation. For spillover, human and livestock are major sources of environmental contamination with a bacterium that is capable of causing disease in diverse host species, and often resistant to antibiotics. Persistent carriage or infection of novel emergent strains of *S. aureus* in wild animal species may pose a public health risk to human and livestock health (13-16). To distinguish between these two possibilities, genomic data from isolates from wild animal populations can be used to infer transmission dates and routes, and to identify genetic changes that may represent host-specific adaptation (5, 17-25).

*S. aureus*, including methicillin resistant *S. aureus* (MRSA), is frequently reported in fish and fishery products, with prevalence ranging from 2-60% (26). The presence of *S. aureus* in fish is concerning because the ingestion of staphylococcal enterotoxins produced by *S. aureus*, can cause food poisoning (27,28). It is widely assumed that S. *aureus* in fish products is a result of contamination during handling and processing (29). However, the combined use of antimicrobial drugs in aquaculture and subsequent contamination of aquatic environments, could contribute to the selection, emergence and spread of antibiotic resistant *S. aureus* (30,31). Supporting this conjecture, *S. aureus* has also been found in fish sampled directly from aquaculture, where fish processing and handling is limited and MRSA have been identified in cage-cultured tilapia in Malaysia (32), tank cultured dusky kob in South Africa (33), and farmed fish in Iran (34). Only one study by Salgueiro *et al*. (2020), performed genome sequencing and found CC398 methicillin-sensitive *S. aureus* (MSSA) (35), most likely human-associated (22,36,37). This study suggests that *S. aureus* is present in fish due to spillover from human populations but does not have the discrimination to test whether there is adaptation and persistence of *S. aureus* within fish populations or aquatic environments.

In this study, we investigate whether *S. aureus* is present in fish from potential source populations in the Scottish Highlands. The Scottish Highlands are among the least densely populated regions of Europe at an average density of 8 persons per km^2^ (38). We sample wild brown trout (*Salmo trutta* L.1758) and their environment, including water and sediment. Brown trout vary in habitat, ranging from highland streams to arctic fjords (39), which allows us to investigate the prevalence of *S. aureus* in areas with varied levels of exposure to known hosts of *S. aureus*. We sample from lochs close to livestock, lochs within bird roosting/feeding areas, sea sites close to human populations, and remote lochs that are isolated from human, bird, and livestock populations. While we readily detected *S. aureus* from multiple organs of a farmed fish in London, we did not detect *S. aureus* in any Scottish trout, nor their environment. These results support the presence of *S. aureus* in farmed fish being due to recent spillover from other host populations.

## Methods

### Calibration of *S. aureus* isolation from fish

To optimise our method of isolation of *S. aureus* from fish, a pilot experiment was carried out on a single rainbow trout (*Oncorhynchus mykiss*) purchased from a fish farm in London, England. Wearing gloves, we swabbed the mouth, vent, gill, and skin with charcoal swabs (Medical Wire Transwabs®), and then inoculated into 10ml of 6% NaCl *Staphylococcus* selective media (A&E Laboratories). We dissected and swabbed the intestine, gill, skin, and heart, and transferred each organ to a sealed processing bag containing 10ml 6% NaCl *Staphylococcus*-selective media, homogenising with a Stomacher (Stomacher80 Laboratory System, Seward Ltd, UK). We took nasal swabs from both researchers carrying out sample processing to control for potential contamination, processed in the same way as fish swabs. We incubated all samples at 37°C in universal tubes for 24h. After incubation, we plated 10-100μL of each culture (depending on the culture cloudiness) onto Brilliance Staph 24 Agar plates (Oxoid, UK) and incubated at 37°C for 24h. For each positive plate, we confirmed *S. aureus* by selecting three colonies for a *femB* PCR (see PCR protocol below).

### Polymerase Chain Reaction (PCR) to amplify *femB*

To confirm the presence of *S. aureus*, we used a colony PCR with *femB* primers. The primers were FemB1 (5′-CAT GGT TAC GAG CAT CAT GG) and FemB2 (5′-AAC GCC AGA AGC AAG GTT TA), leading to an *S. aureus*-specific 447bp PCR product (41). We touched a pipette tip on a single *S. aureus* colony and mixed it with 20 μl of water in a PCR tube. We boiled the samples for 5 min at 95ᵒC. Each sample contained 2μl of boiled cell solution and 18μl of the PCR master mix with MyTaq DNA Polymerase (Bioline). The PCR cycling conditions were 95 ᵒC 5 min, 30 cycles (95 ᵒC-15 s, 58ᵒC – 10s, 72ᵒC 30s), 72ᵒC for 10 min. We added 2.5μL Sybr® Safe DNA gel stain (Thermo Fisher, UK) to 100μL agarose solution, which was 1% w/v agarose dissolved by heating in a 1×TBE buffer (Tris-borate-EDTA). We loaded 15μL of PCR reaction mixture onto each well, with a 5μL 5 HyperLadderTM 100bp (Bioline) to confirm product size. Electrophoresis was performed in a Sigma-Aldrich electrophoresis tank with 1 ×TBE at 100V for 40 minutes. Electrophoresed gels were visualised under blue light, and their images visualised with the GelDocTM XR System Imager (BioRad).

### Sequencing

For all confirmed and suspected *S. aureus* positive samples from the farmed fish, two colonies were selected for each sampling method (swabbing and tissue samples) per positive sampling site (gill, intestine, skin). Genomic DNA was extracted from overnight cultures grown in Tryptic Soy Broth (TSB) at 37°C with 200rpm shaking using the MasterPure Gram Positive DNA Purification Kit (Cambio, UK). Illumina library preparation and Hi-Seq sequencing were carried out as described in (41).

### Genome assembly and MLST typing

Published genome sequence data from isolates of CC45 were downloaded from the European Nucleotide Archive (ENA) and subsampled to represent host and geographical diversity (Table S1). Sequence data from both newly sequenced and publicly available CC45 isolates were assembled using Spades v.3.12.0 (42). Adapters and low-quality reads were removed with Cutadapt v1.16 (43) and Sickle v1.33 (44) and screened for contamination using FastQ Screen v0.12.0 (45). We identified optimal k-mers based on average read lengths for each genome. All *de novo* assemblies were evaluated using QUAST v.5.0.1 (46) and reads were mapped back to *de novo* assemblies to investigate polymorphism (indicative of mixed cultures) using Bowtie2 v1.2.2 (47). All assemblies were of good quality (i.e., N50 <10,000, contigs smaller than 1kb contributing to >15% of the total assembly length, total assembly length outside of the median sequence length +/-one standard deviation, or >1500 polymorphic sites).

### Phylogenetic analyses and genome annotation

Reference-mapped assemblies were generated using Bowtie2 v1.2.2, using the reference genome LGA251 (GenBank accession no. GCA_000237265.1) (47). All genomes had average coverage <50x or with >10% missing sites. Recombination was identified in the reference-mapped alignment using Gubbins v2.3.1 and recombinant sites were masked from phylogenetic analyses (48). Sites that had either recombination detected or missing data were excluded from phylogenetic analyses. Phylogenetic reconstruction was carried out for the reference-mapped alignment with RAxML (v8.2.4) using the GTR+Γ model and 1,000 bootstraps (49) and rooted using an isolate from CC398 (SRR445234). To investigate the diversity among closely related isolates, we examined the reference mapped alignment to identify single nucleotide polymorphisms (SNPs). The alignment after recombination stripping was uploaded to Geneious 2020.0.4 (https://www.geneious.com), and all regions that were within 100bp of large regions containing missing and regions containing SNP clusters > 5 SNPs located within 50bp were removed. Next, the SNPs were extracted, and an MSTree V2 was constructed using GrapeTree (50).

We identified antibiotic resistance genes using the *Pathogenwatch* AMR prediction module (Wellcome Sanger Institute), which uses *BLASTn* (51) with a cut-off of 75% coverage and 80–90% identity threshold (depending on the gene) against a *S. aureus* antimicrobial resistance database. Presence of φSa3 prophages was established through searching for genes in the human immune evasion gene cluster using *BLASTn* (51) with a cut-off of 90% coverage and 90% identity threshold. The query human immune evasion genes were extracted from a reference genome (assembly accession: GCA_900324385.1). Presence of enterotoxins was investigated through searching for relevant proteins from a database compiled by Merda and Felten *et al*., 2020, using *tBLASTn* cut-off of 80% coverage and 80% (52).

### Sample size and statistical analysis

To guide sample size design, we calculated the probability of detecting one or more positive sample with varying levels of prevalence using the *pbinom* function within the software package R (53). We show that 30 fish samples give us a 95% probability of detecting a positive sample if the true prevalence is around 10% (Figure S1). Prevalence of *S. aureus* in fish varies but is typically greater than 10% (26, 33). Therefore, we chose to sample 30 fish from each of our four habitats. In total, 120 fish gives a 95% probability of detecting a positive if the true prevalence at all sites is around 3%. We calculated 95% confidence intervals shown in Table 1 on our observed prevalence, using the *binom*.*test* function in R (53).

**Table 1:** Prevalence of *S. aureus* in the environment and from brown trout in different habitats.

| Habitat | No. of sampled lochs/sites | No. environment samples | No. of positive environmental samples | No. trout samples | No. of positive isolates | Prevalence in trout |
| --- | --- | --- | --- | --- | --- | --- |
| All habitats | 16 | 32 | 0 | 123 | 0 | 0 (0-0.03) |
| Bird Lochs | 4 | 8 | 0 | 42 | 0 | 0 (0-0.08) |
| Isolated Lochs | 5 | 10 | 0 | 48 | 0 | 0 (0-0.07) |
| Livestock Lochs | 4 | 8 | 0 | 30 | 0 | 0 (0-0.12) |
| Human (Sea) | 3 | 6 | 0 | 3 | 0 | 0 (0-0.71) |

### Fish and environmental sampling

All sampling was carried out in July 2019. Brown trout were captured by fly-fishing and transferred, along with fresh loch/sea water, to a sterilised bucket. We swabbed the vents and gills whilst the trout was alive, with operators wearing sterile gloves to prevent cross-contamination. Fish larger than 250mm were released (a condition of the landowners). We euthanized fish smaller than 250mm via a blow to the back of the head with a sterilised priest, placed them in a specimen bag and stored them in an icebox until later transferring to a -80°C freezer, for long term storage. The sites were selected based on their isolation from (Isolated Lochs) or their proximity (<1 km) to human frequented sites, including camping sites (Human), livestock grazing sites (Livestock Lochs) and bird nesting sites (Bird Lochs). All habitat types, except for Bird Lochs, were represented by either two or three site clusters >21 km apart.

To test whether *S. aureus* was present in the environment, we collected water and sediment samples from 25 sites, including all sites where we sampled trout. Sampling was carried out with operators wearing sterile gloves to prevent cross-contamination. We took water (3.5 litres) from approximately 5-10 cm depth using sterile 1000ml and 500ml containers. We collected sediment (approximately 50g per location) to sterile containers using a sterilised plastic scoop from littoral areas that were a) > 150mm below the water surface; and b) undisturbed. We kept both water and sediment samples chilled at 4°C, until analysis. Samples were processed immediately upon arrival at the laboratory. We took nasal swabs from both researchers carrying out sample processing to control for potential contamination.

### *S. aureus* isolation from environmental samples

To isolate *S. aureus* from water, we filtered three aliquots of 1000 ml per water sample through 0.45μm 47mm white gridded mixed ester cellulose membranes (Merck, USA). Membranes were placed on 55 mm Baird-Parker with potassium tellurite enrichment agar plates and incubated at 37°C for 48 hours. To isolate *S. aureus* from sediment, we transferred three 10 g (wet weight) sediment samples (per location) to sterile 50 ml centrifuge tubes. We added 25 ml of 2x concentrated Baird-Parker with potassium tellurite enrichment medium to each sample. After vortexing for approximately 45 s, the supernatant from each sample was transferred to a new sterile tube and incubated until a black precipitate formed (up to seven days). After this, the samples were vortexed for 30s, and 100μl was plated on 100 mm Baird-Parker with potassium tellurite enrichment agar plates and incubated at 37°C for 48 hours. For both water and sediment samples, after incubation, we transferred three presumptive *S. aureus* colonies (black colonies) onto fresh Brilliance Staph 24 plates (Oxoid, UK) and incubated at 37°C for 24 hours. For each potential positive plate, we confirmed *S. aureus* by selecting three colonies for a *femB* PCR (see PCR protocol below).

### Antibiotic susceptibility testing

Antibiotic susceptibility testing was carried out with the Vitek 2 system (AES software, bioMérieux, Marcy l’Étoile, France) according to the manufacturer’s instructions. The isolates were plated onto Columbia Blood Agar (Oxoid Deutschland) and incubated at 37°C for 18-24h. One colony was picked with a sterile swab and mixed in the saline by vortexing. The OD_600_ .was measured and adjusted to 0.5-0.63. The inoculum was prepared by transferring 280μl to 3ml saline. Next, each sample was loaded into a VITEK® 2 AST-GP80 card. Susceptibility cards were interpreted according to the Clinical and Laboratory Standards Institute (CLSI) breakpoints (54).

## Results

### *S. aureus* sample calibration and whole genome diversity from a farmed fish

To establish the most effective method for detecting *S. aureus* from fish, we swabbed and homogenised tissue from different body sites of a farmed Rainbow trout (*Oncorhynchus mykiss*). *S. aureus* was successfully isolated from all sites, apart from a heart tissue sample and a mouth swab (Table 2). All positive tissue samples were also detected using charcoal swabs, suggesting that tissue processing is unnecessary for *S. aureus* detection from positive sites. Both swabs from researchers carrying out the sample processing were negative for *S. aureus*.

**Table 2:**
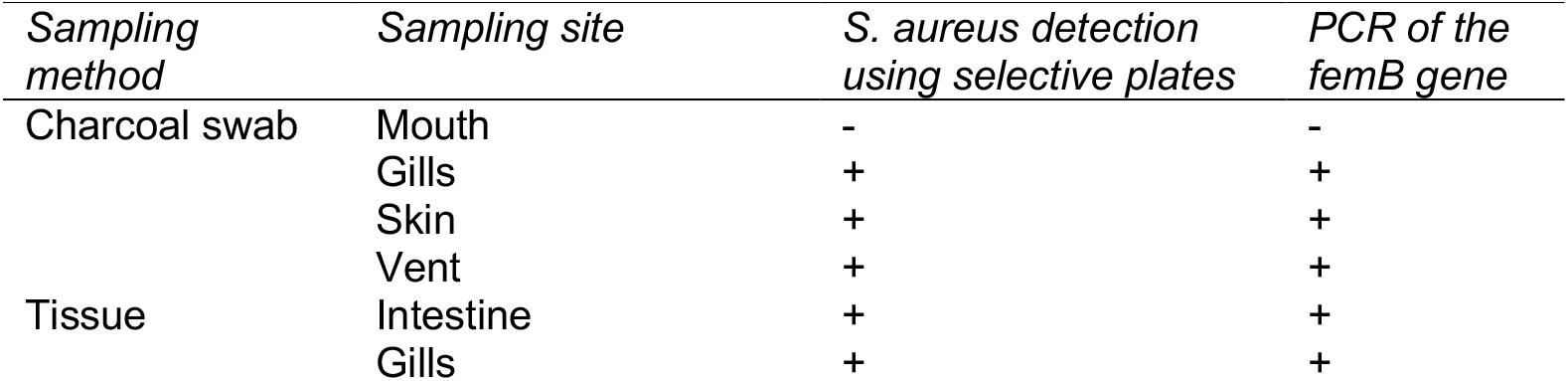
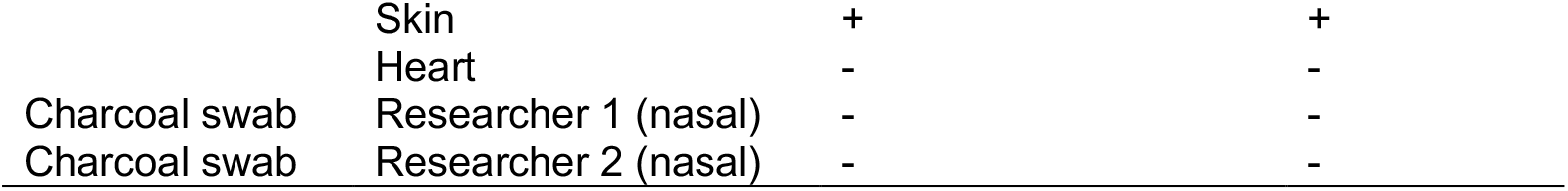
The detection of *S. aureus* from a single rainbow trout purchased from a fish farm in London, England.

To investigate within fish diversity, whole-genome sequencing was carried out for 12 *S. aureus* isolated from the different body sites. All isolates were identified as a strain type (ST) 54, which is part of clonal complex (CC) 45. Figure 1 shows a grape plot from a core genome alignment mapped to the *S. aureus* reference genome, LGA251. The isolates are not clustered by sampling site (intestine, gill, and skin) or by sampling method (charcoal swabbing and tissue). The isolates are very closely related, with the maximum distance between two isolates being 12 single nucleotide polymorphisms (SNPs). Based on an SNP clock rate of ∼3.5 SNPs/genome/year in ST22 (55), we estimated that these isolates’ most recent common ancestor would have dated to around 2017 (two years before sampling).

**Figure 1.**
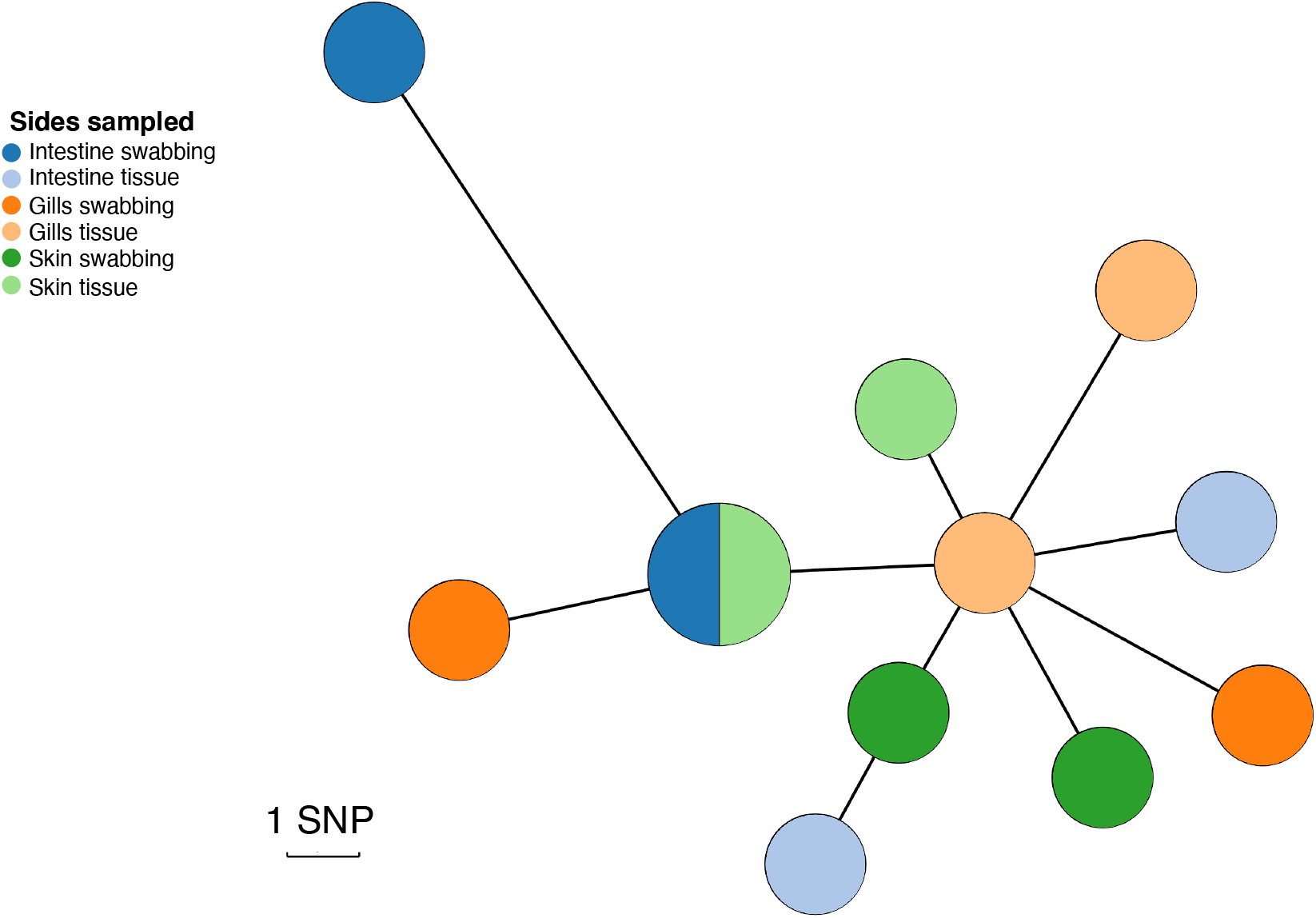
The genetic diversity of *S. aureus* isolated from three different anatomical sites in a single farmed rainbow trout from a London fish farm. A minimum-spanning tree was constructed using an alignment of the 12 fish isolates, created by mapping to the *S. aureus* reference genome, LGA251. Points represent groups of identical isolates, with point size correlated with the number isolates. Due to the low number of SNPs, each SNP was manually checked to ensure that they were not a consequence of mapping error. The colours represent different sampling fish sites: intestine (blue), gills (orange), and skin (green). Isolates extracted using charcoal swabs are indicated in a darker shade, where the tissue samples are indicated in a lighter shade.

To place the farmed rainbow trout isolates within the known diversity of CC45, we assembled the genomes from publicly available sequencing data from a set of CC45 isolates (Table S1). CC45 was mostly associated with human hosts (71%) but also identified in non-human hosts. The phylogeny of CC45 shows that all of the samples we collected from the farmed trout fall in a single clade, with a human *S. aureus* from the UK being the closest relative (Figure 2a and 2b). The mean SNP distance between this closest relative and the rainbow trout clade is 165 SNPs. Assuming the same clock rate as above, we estimate that the fish clade diverged from the human isolate around 1995. We identified genes carried on a φSa3 prophage that is known to be associated with human immune evasion in most isolates in our sample of CC45, including those from the farmed rainbow trout (Figure 2d). This suggests that this CC is adapted to the human host, and that its presence in other species is a consequence of recent spillover.

**Figure 2:**
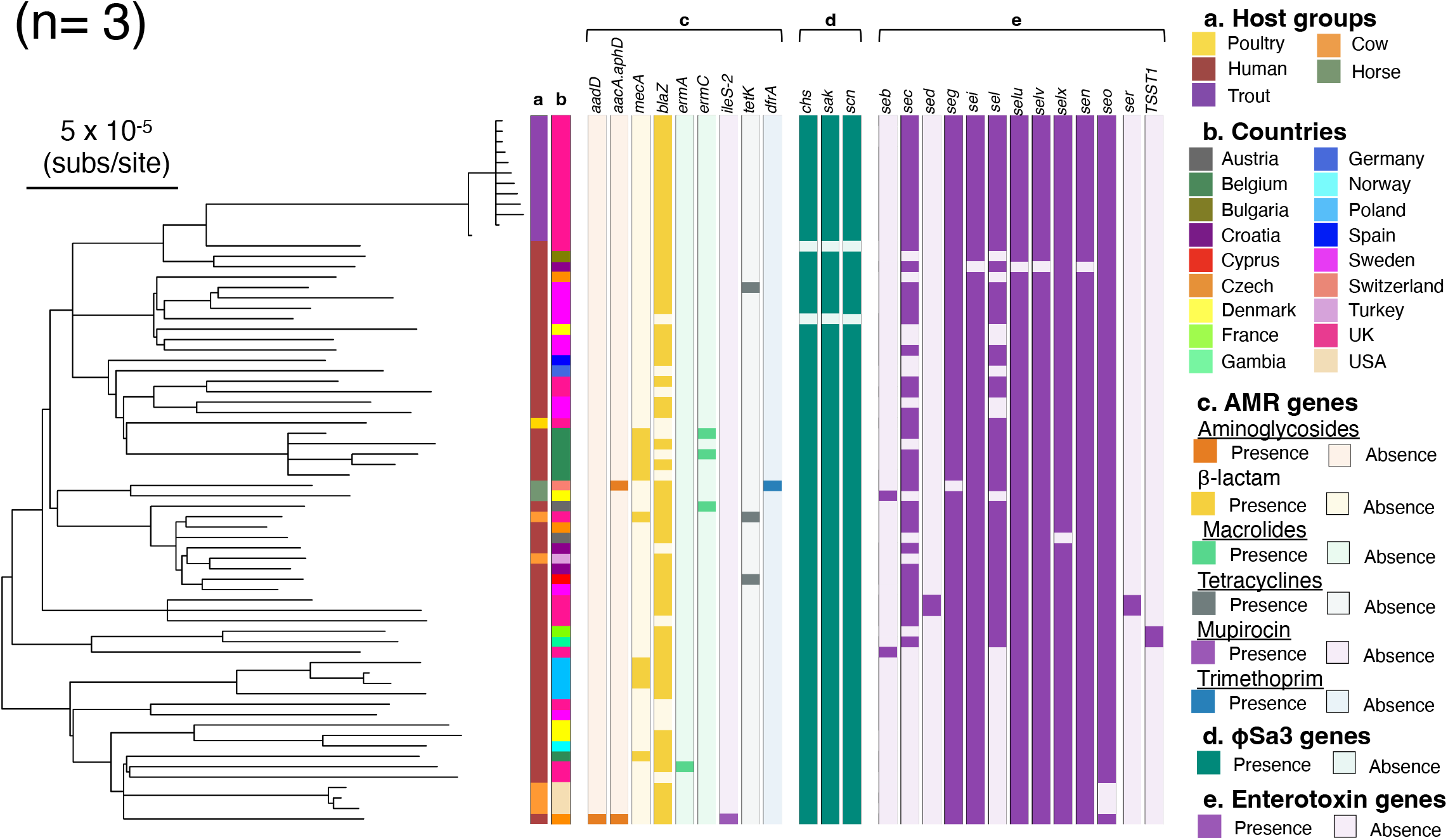
*S. aureus* isolates from a single rainbow trout purchased from a fish farm in London, England fall within the diversity of CC45. A maximum-likelihood phylogeny of 68 isolates of CC45 based on an LGA251 reference-mapped alignment with recombination removed and rooted using an outgroup from CC398 (with branches <70 bootstrap support collapsed). Outer rings describe (a) the host groups isolates were sampled from, (b) country, presence and absence of (c) AMR genes, (d) φSa3 prophage functional genes, and I enterotoxin genes.

CC45 is, in general, methicillin-sensitive (56) and only 2/68 of the isolates in our sample carry the gene associated with methicillin resistance (*mecA*). Genes associated with resistance to other antibiotics are also rare in this CC. Consistent with the low levels of resistance in this clade the isolates from the farmed trout were found to be methicillin-sensitive and carry no resistance genes except for the *blaZ* gene that confers resistance to benzylpenicillin (Figure 2c). Phenotypic resistance was confirmed by selecting two representative isolates, which both matched the genotypic data (Table S2).

We investigated the potential of CC45 isolates to cause Staphylococcal food poisoning by checking for the presence of staphylococcal enterotoxins. The isolates we sampled from the farmed fish carry nine enterotoxin coding genes, which are common in CC45.

### Absence of *S. aureus* in wild population of brown trout

We sampled fish from four habitat types that are likely to show variation in spillover exposure. These are habitats with exposure to (1) human, (2) livestock, and (3) avian hosts, as well as (4) very isolated sampling sites (Figure 3). We aimed to sample 30 fish from each of our habitats and collect a total number of 120 fish (see methods). While our eventual sample size was 123, we were unable to reach the desired number of trout samples from the sea (exposure to human populations). We sampled 123 brown trout from 16 sites from four habitats (between 1-11 fish from each loch). We collected water and sediment samples from 23 sites from four habitats and three additional sites not categorised into any of the habitats (Tables S3-S5). All researchers carrying out fishing and sampling were swabbed and found to be negative for *S. aureus*.

**Figure 3.**
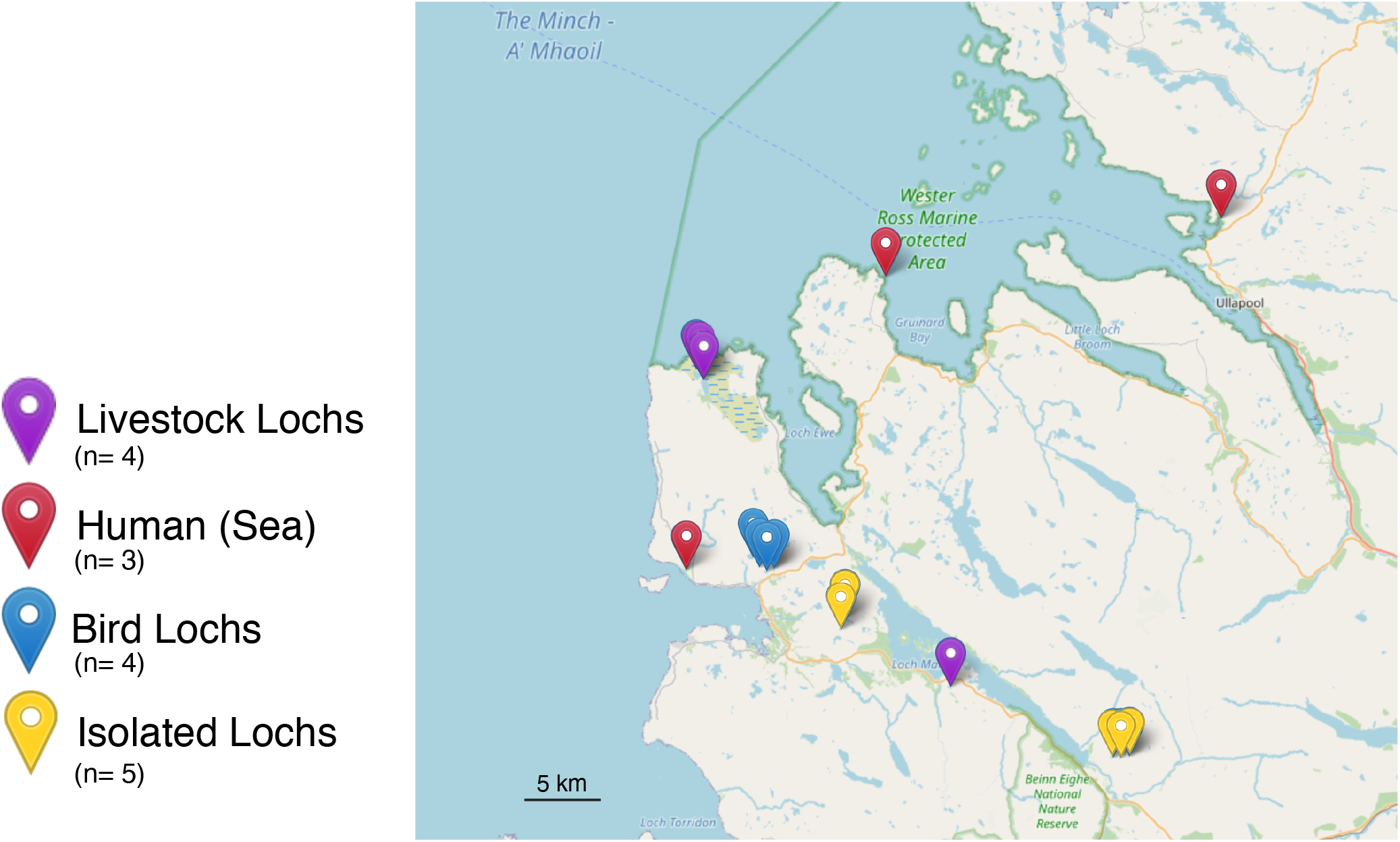
Location of all the sampling sites in Wester Ross, Scotland. The sites include bird, isolated and livestock lochs and sea. The map was constructed using R (53) Leaflet library (v.2.0.4.1).

We did not detect any positive samples in any of the 123 wild brown trout, nor in their environment (Table 1). Overall, these results suggest that *S. aureus* is either absent or present at very low prevalence in Scottish lochs.

## Discussion

Our study did not detect *S. aureus* in wild populations of brown trout, nor in the Scottish lochs that they inhabit. All sites with different (although all relatively low) levels of exposures to species that are known hosts of *S. aureus*, including livestock, humans and birds, were negative for *S. aureus*. This suggests the absence or a very low prevalence of *S. aureus* in these populations. Our results are consistent with another study testing for the presence of *S. aureus* from different species of 168 wild fish caught by trawling and rod fishing in sea populations from Japan, where no *S. aureus* was detected (57). Overall, these results argue that the presence of *S. aureus* in farmed fish and fish products is not due to the adaptation of *S. aureus* to wild fish populations.

Previous studies have documented the presence of *S. aureus*, including MRSA, in fish farms and in fish products (32,33). We detected *S. aureus* in both internal and external organs of a single farmed fish using aseptic techniques, suggesting that its presence is not due to contamination via handling. *S. aureus* could be adapted to and persisting within the fish host. However, the *S. aureus* genomic diversity within the single fish is consistent with a date of acquisition (i.e., approximately two years) that is likely older than the lifespan of the fish (the fish weight was 300g, which is consistent with an age of less than eight months (58)). Given the lack of clustering by anatomical location, it is more likely that the fish acquired *S. aureus* through passive filtration of its external environment. Previous studies suggested that MRSA can also survive for extended periods of time in the sea and river, and marine fresh water (59,60).

*S. aureus* may be adapted to the conditions within aquaculture. We detected the *blaZ* gene in the fish farm isolates. Β-lactamase production (encoded by *blaZ* gene) renders *S. aureus* resistant to benzylpenicillin, phenoxymethylpenicillin, ampicillin, amoxicillin, piperacillin and ticarcillin (61). Amoxicillin and ampicillin are commonly used in aquaculture worldwide (62,63). Therefore, the *blaZ* gene could provide *S. aureus* with a selective advantage over other non-resistant bacteria within this environment. However, our results show that the fish farm rainbow trout isolates are part of CC45, which is associated with both nasal colonization and bloodstream infections in humans (56). The fish farm isolates nest within a more diverse clade of human isolates and contains a φSa3 prophage, which encodes a human immune evasion cluster (64). Our results therefore suggest that the presence of *S. aureus* in this aquaculture environment is due to recent spillover from human populations. This is consistent with another study that showed the presence of a CC398 MSSA isolate from a gilthead seabream (35), which most likely originated from a human-associated lineage of CC398 (22,36,37). *S. aureus* have been recovered from municipal (65–68), hospital (69), and agricultural wastewaters/sewage (70), representing potential sources of human environmental contamination.

We detected nine enterotoxin genes in all our farmed fish isolates. Staphylococcal enterotoxins have also been detected in other studies of *S. aureus* in fish and fishery products (71,72). Staphylococcal food poisoning is caused by the ingestion of any of the 27 characterised Staphylococci enterotoxins (27,28). The toxins can have neurotic or superantigenic activity which result in vomiting or fever respectively (73). The Staphylococci enterotoxins have high tolerance to denaturing conditions, such as low pH or heat and even when ingested in low quantities, can be toxic to humans (74). This suggests that fish from aquaculture could constitute a risk of food poisoning (73,74).

This study also optimized a sampling approach using farmed fish to calibrate a *S. aureus* isolation protocol. Previous techniques for *S. aureus* isolation relied on the fish being euthanized, whereas this approach allowed fish to be returned to their environment after sampling (32,75,76).

The absence of *S. aureus* in wild fish populations combined with whole genome sequencing from a farmed fish, suggest that the presence of *S. aureus* in fish is the result of spillover from source populations. Nevertheless, wider whole genome sequencing of *S. aureus* from aquaculture, including the environment and fish farm workers, is needed to determine the origin and mechanisms of persistence of *S. aureus* in this environment. Fish and seafood consumption is predicted to increase by 27% by 2030 and much of this increase will be supplied by growth in the aquaculture sector (77). Aquaculture is known to introduce and amplify new pathogens (78), and it relies heavily on antibiotics to combat infectious diseases (30,79,80). While our study found no evidence of adaptation of *S. aureus* to fish, growth in the size and density of fish farms could create more opportunities for *S. aureus* to adapt to aquaculture and to fish and could also promote the evolution and spread of strains that are resistant to antibiotics. This would have potential impacts on both food security and human health.

## Acknowledgements

The authors acknowledge the assistance of Wester Ross Fisheries Trust without whose expertise and resources this study would not have been possible. The authors acknowledge the comments on an early draft of Camille Bonneaud and Julian Parkhill. MM was funded by the Medical Research Council, co-funded by the Raymond and Beverly Sackler Fund. GGRM and LAW were supported by a Sir Henry Dale Fellowship jointly funded by the Wellcome Trust and the Royal Society (109385/Z/15/Z). GGRM was also supported by a ZELS BBSRC award (BB/L018934/1) and a Research Fellowship at Newnham College. AD was funded by the Engineering and Physical Sciences Research Council (EP/P029426/1).

---

**Figure S1:**
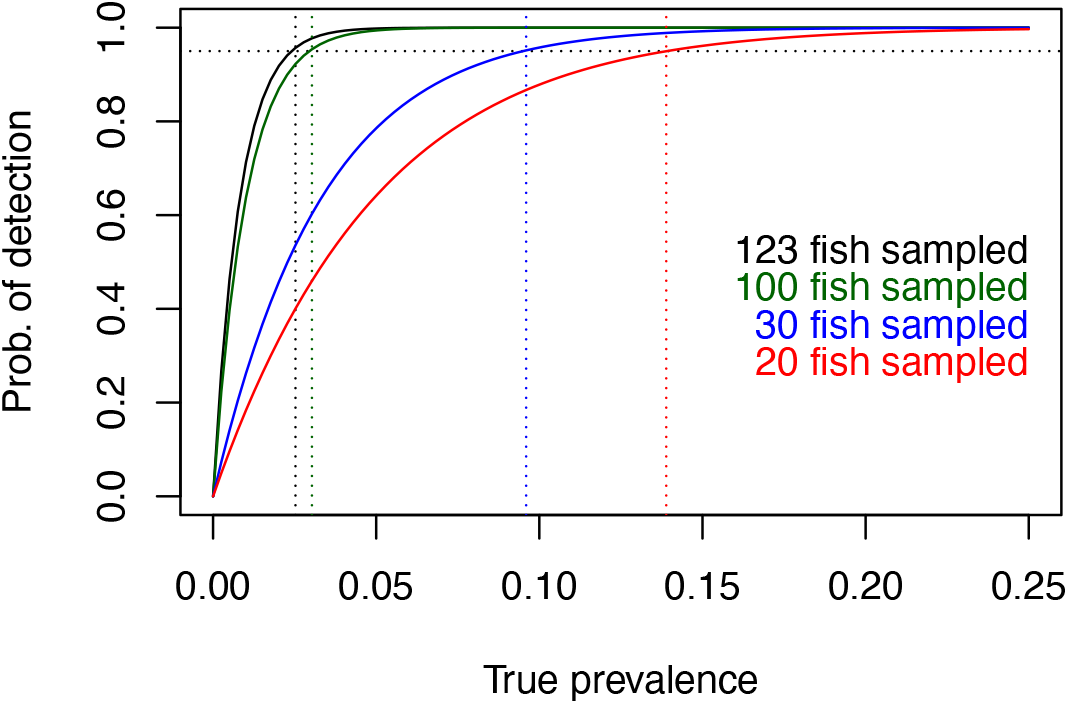
The probability of detecting one or more positive *S. aureus* isolate from fish samples (y axis), given differences in the true prevalence (x axis). The coloured lines correspond to different numbers of fish samples. For example, 20 fish samples gives a 95% chance of detecting one or more positive *S. aureus* isolate if the true prevalence is around 0.14.

## References

1. O’Gara JP. 2017. Into the storm: Chasing the opportunistic pathogen Staphylococcus aureus from skin colonisation to life-threatening infections. Environ Microbiol 19(10):3823–3833.

2. Loeffler A, Pfeiffer DU, Lindsay JA, Magalhães RJS, Lloyd DH. 2011. Prevalence of and risk factors for MRSA carriage in companion animals: A survey of dogs, cats and horses. Epidemiol Infect 139(7):1019–28.

3. Harrison EM, Weinert LA, Holden MTG, Welch JJ, Wilson K, Morgan FJE, Harris SR, Loeffler A, Boag AK, Peacock SJ, Paterson GK, Waller AS, Parkhill J, Holmes MA. 2014. A shared population of epidemic methicillin-resistant Staphylococcus aureus 15 circulates in humans and companion animals. Mbio 5(3).

4. Peton V, Le Loir Y. 2014. Staphylococcus aureus in veterinary medicine. Infect Genet Evol 21:602–615.

5. Lowder BV, Guinane CM, Zakour NLB, Weinert LA, Conway-Morris A, Cartwright RA, Simpson AJ, Rambaut A, Nübel U, Fitzgerald JR. 2009. Recent human-to-poultry host jump, adaptation, and pandemic spread of Staphylococcus aureus. Proc Natl Acad Sci U S A 106(46):19545–50.

6. Fitzgerald JR. 2012. Livestock-associated Staphylococcus aureus: Origin, evolution and public health threat. Trends Microbiol 20(4):192–198.

7. Mrochen DM, Schulz D, Fischer S, Jeske K, El Gohary H, Reil D, Imholt C, Trübe P, Suchomel J, Tricaud E, Jacob J, Heroldová M, Bröker BM, Strommenger B, Walther B, Ulrich RG, Holtfreter S. 2018. Wild rodents and shrews are natural hosts of Staphylococcus aureus. Int J Med Microbiol 308(6):590–597.

8. Schaumburg F, Alabi AS, Köck R, Mellmann A, Kremsner PG, Boesch C, Becker K, Leendertz FH, Peters G. 2012. Highly divergent Staphylococcus aureus isolates from African non-human primates. Environ Microbiol Rep 4(1):141–146.

9. Schaumburg F, Mugisha L, Peck B, Becker K, Gillespie TR, Peters G, Leendertz FH. 2012. Drug-resistant human Staphylococcus aureus in sanctuary apes pose a threat to endangered wild ape populations. Am J Primatol 74(12):1071–1075.

10. Gómez P, Lozano C, Camacho MC, Lima-Barbero JF, Hernández JM, Zarazaga M, Höfle Ú, Torres C. 2016. Detection of MRSA ST3061-t843-mecC and ST398-t011-mecA in white stork nestlings exposed to human residues. J Antimicrob Chemother 71(1):53–57.

11. Monecke S, Gavier-Widén D, Hotzel H, Peters M, Guenther S, Lazaris A, Loncaric I, Müller E, Reissig A, Ruppelt-Lorz A, Shore AC, Walter B, Coleman DC, Coleman R. 2016. Diversity of Staphylococcus aureus Isolates in European Wildlife. PloS One 11(12):e0168433.

12. Centers for Disease Control and Prevention (CDC). 2009. Methicillin-Resistant Staphylococcus aureus skin infections from an elephant calf – San Diego, California, 2008. MMWR Morb Mortal Wkly Rep 58(8):194–198.

13. Kreuder Johnson C, Hitchens PL, Smiley Evans T, Goldstein T, Thomas K, Clements A, Joly DO, Wolfe ND, Daszak P, Karesh WB, Mazet JK. 2015. Spillover and pandemic properties of zoonotic viruses with high host plasticity. Sci Rep. 5(1):e14830.

14. Morse SS, Mazet JA, Woolhouse M, Parrish CR, Carroll D, Karesh WB, Zambrana-Torrelio C, Lipkin WI, Daszak P. 2012. Prediction and prevention of the next pandemic zoonosis. Lancet 380(9857):1956–1965.

15. Sharp PM, Hahn BH. 2011. Origins of HIV and the AIDS pandemic. Cold Spring Harb Perspect Med 1(1):a006841.

16. Fitzgerald RJ, Holden MTG. 2016. Genomics of Natural Populations of Staphylococcus aureus. Annu Rev Microbiol 70:459–478.

17. Guinane CM, Zakour NLB, Tormo-Mas MA, Weinert LA, Lowder B V, Cartwright RA, Smyth DS, Smyth CJ, Lindsay JA, Gould KA, Witney A, Hinds J, Bollback JP, Rambaut A, Penadés JR, Fitzgerald JR. 2010. Evolutionary genomics of Staphylococcus aureus reveals insights into the origin and molecular basis of ruminant host adaptation. Genome Biol Evol 2:454–466.

18. De Jong NWM, Vrieling M, Garcia BL, Koop G, Brettmann M, Aerts PC, Ruyken M, van Strijp JAG, Holmes MA, Harrison EM, Geisbrecht BV, Rooijakkers SHM. 2018. Identification of a staphylococcal complement inhibitor with broad host specificity in equid Staphylococcus aureus strains. J Biol Chem 293(12):4468–4477.

19. Koop G, Vrieling M, Storisteanu DML, Lok LSC, Monie T, Van Wigcheren G, Raisen C, Ba X, Gleadall N, Hadjirin N, Timmerman AJ, Wagenaar JA, Klunder HM, Fitzgerald JR, Zadoks R, Paterson GK, Torres C, Waller AS, Loeffler A, Loncaric I, Hoet AE, Bergström K, De Martino L, Pomba C, de Lencastre H, Slama KB, Gharsa H, Richardson EJ, Chilvers ER, de Haas C, van Kessel K, van Strijp JAG, Harrison EM, Holmes MA. 2017. Identification of LukPQ, a novel, equid-adapted Leucocidin of Staphylococcus aureus. Sci Rep 7:e40660.

20. Rooijakkers SHM, Van Wamel WJB, Ruyken M, Van Kessel KPM, Van Strijp Jag. 2005. Anti-opsonic properties of staphylokinase. Microbes Infect 7(3):476–484.

21. Van Wamel WJB, Rooijakkers SHM, Ruyken M, Van Kessel KPM, Van Strijp Jag. 2006. The innate immune modulators staphylococcal complement inhibitor and chemotaxis inhibitory protein of Staphylococcus aureus are located on β-hemolysin-converting bacteriophages. J Bacteriol 188(4):1310–1315.

22. Price LB, Stegger M, Hasman H, Aziz M, Larsen J, Andersen PS, Pearson T, Waters AE, Foster JT, Schupp J, Gillece J, Driebe E, Liu CM, Springer B, Zdovc I, Battisti A, Franco A, Zmudzki J, Schwarz S, Butaye P, Jouy E, Pomba C, Porrero MC, Ruimy R, Smith TC, Robinson DA, Weese JS, Arriola CS, Yu F, Laurent F, Keim P, Skov R, Aarestrup FM. 2012. Staphylococcus aureus CC398: Host adaptation and emergence of methicillin resistance in livestock. mBio 3:1–6.

23. Resch G, François P, Morisset D, Stojanov M, Bonetti EJ, Schrenzel J, Sakwinska O, Moreillon P. 2013. Human-to-Bovine Jump of Staphylococcus aureus CC8 Is Associated with the Loss of a β-Hemolysin Converting Prophage and the Acquisition of a New Staphylococcal Cassette Chromosome. PloS One 8(3):e58187.

24. Spoor LE, McAdam PR, Weinert LA, Rambaut A, Hasman H, Aarestrup FM, Kearns AM, Larsen AR, Skov RL, Fitzgerald JR. 2013. Livestock origin for a human pandemic clone of community-associated methicillin-resistant Staphylococcus aureus. mBio 4(4):e00356–13.

25. Richardson J, Bacigalupe R, Harrison EM, Weinert LA, Richardson EJ, Lycett S, Vrieling M, Robb K, Hoskisson PA, Holden MTG, Feil EJ, Paterson GK, Tong SYC, Shittu A, van Wamel W, Aanensen DM, Parkhill J, Peacock SJ, Corander J, Holmes MA, Fitzgerald JR. 2018. Gene exchange drives the ecological success of a multi-host bacterial pathogen. Nat Ecol Evol 2:1468–1478.

26. Vaiyapuri M, Joseph TC, Rao BM, Lalitha KV, Prasad MM. 2019. Methicillin-Resistant Staphylococcus aureus in Seafood: Prevalence, Laboratory Detection, Clonal Nature, and Control in Seafood Chain. J Food Sci 84(12):3341–3351.

27. Tirado C, Schmidt K. 2001. WHO surveillance programme for control of foodborne infections and intoxications: Preliminary results and trends across greater Europe. J Infect 43(1):80–84.

28. Hennekinne JA, De Buyser ML, Dragacci S. 2012. Staphylococcus aureus and its food poisoning toxins: Characterization and outbreak investigation. FEMS Microbiol Rev 36(4):815–836.

29. Todd ECD, Greig JD, Bartleson CA, Michaels BS. 2008. Outbreaks where food workers have been implicated in the spread of foodborne disease. Part 4. Infective doses and pathogen carriage. J Food Prot 71(11):2339–2373.

30. Schar D, Sommanustweechai A, Laxminarayan R, Tangcharoensathien V. 2018. Surveillance of antimicrobial consumption in animal production sectors of low- and middle-income countries: Optimizing use and addressing antimicrobial resistance. PloS Med 15(3) :e1002521.

31. Van Boeckel TP, Brower C, Gilbert M, Grenfell BT, Levin SA, Robinson TP, Teillant A, Laxminarayan R. 2015. Global trends in antimicrobial use in food animals. Proc Natl Acad Sci U S A 112(18):5649–5654.

32. Atyah MAS, Zamri-Saad M, Siti-Zahrah A. 2010. First report of methicillin-resistant Staphylococcus aureus from cage-cultured tilapia (Oreochromis niloticus). Vet Microbiol 144(3–4):502–504.

33. Fri J, Ndip RN, Njom HA, Clarke AM. 2017. First report of methicillin-resistant Staphylococcus aureus in tank cultured dusky kob (Argyrosomus japonicus), and evaluation of three phenotypic methods in the detection of MRSA. J Food Saf 38(1):e12411.

34. Arfatahery N, Davoodabadi A, Abedimohtasab T. 2016. Characterization of Toxin Genes and Antimicrobial Susceptibility of Staphylococcus aureus Isolates in Fishery Products in Iran. Sci Rep 6:34216.

35. Salgueiro V, Manageiro V, Bandarra NM, Ferreira E, Clemente L, Caniça M. 2020. Genetic relatedness and diversity of Staphylococcus aureus from different reservoirs: Humans and animals of livestock, poultry, zoo, and aquaculture. Microorganisms 8(9):1345.

36. Gonçalves da Silva A, Baines SL, Carter GP, Heffernan H, French NP, Ren X, Seemann T, Bulach D, Kwong J, Stinear TP, Howden BP, Williamson DA. 2017. A phylogenomic framework for assessing the global emergence and evolution of clonal complex 398 methicillin-resistant Staphylococcus aureus. Microb Genom 3(1):e000105.

37. Matuszewska M, Murray GG, Ba X, Wood R, Holmes MA, Weinert LA. 2022. Stable antibiotic resistance and rapid human adaptation in livestock-associated MRSA. Elife 11:e74819.

38. Geography | Highland profile – key facts and figures | The Highland Council [Internet]. [cited 2022 Aug 17]. Available from: https://www.highland.gov.uk/info/695/council_information_performance_and_statistics/165/highland_profile_-_key_facts_and_figures

39. Ferguson A, Reed TE, Cross TF, McGinnity P, Prodöhl PA. 2019. Anadromy, potamodromy and residency in brown trout Salmo trutta: the role of genes and the environment. J Fish Biol 95(3):692–718.

40. Paterson GK, Larsen AR, Robb A, Edwards GE, Pennycott TW, Foster G, Mot D, Hermans K, Baert K, Peacock SJ, Parkhill J, Parkhill RN, Holmes MA. 2012. The newly described mecA homologue, mecALGA251, is present in methicillin-resistant Staphylococcus aureus isolates from a diverse range of host species. J Antimicrob Chemother 67(12):2809–2813.

41. Harrison EM, Paterson GK, Holden MTG, Larsen J, Stegger M, Larsen AR, Petersen A, Skov RL, Christensen JM, Zeuthen AB, Heltberg O, Harris SR, Zadoks RN, Parkhill J, Peacock SJ, Holmes MA. 2013. Whole genome sequencing identifies zoonotic transmission of MRSA isolates with the novel mecA homologue mecC. EMBO Mol Med 5(4):509–515.

42. Bankevich A, Nurk S, Antipov D, Gurevich AA, Dvorkin M, Kulikov AS, Lesin VM, Nikolenko SI, Pham S, Prjibelski AD, Pyshkin AV, Sirotkin AV, Vyahhi N, Tesler G, Alekseyev MA, Pevzner PA. 2012. SPAdes: A New Genome Assembly Algorithm and Its Applications to Single-Cell Sequencing. J Comput Biol 19(5):455–477.

43. Martin M. 2011. Cutadapt removes adapter sequences from high-throughput sequencing reads. EMBnet J 17(1):10.

44. Joshi N, Fass J. 2011. Sickle: a sliding-window, adaptive, quality-based trimming tool for FastQ files.

45. Wingett SW, Andrews S. 2018. FastQ Screen: A tool for multi-genome mapping and quality control. F1000Research. 7:1338.

46. Gurevich A, Saveliev V, Vyahhi N, Tesler G. 2013. QUAST: quality assessment tool for genome assemblies. Bioinformatics. 29:1072–1075.

47. Langmead B, Salzberg SL. 2012. Fast gapped-read alignment with Bowtie 2. Nat Methods 2012 9:4. 9:357–359.

48. Croucher NJ, Page AJ, Connor TR, Delaney AJ, Keane JA, Bentley SD, Parkhill J, Harris SR. 2015. Rapid phylogenetic analysis of large samples of recombinant bacterial whole genome sequences using Gubbins. Nucleic Acids Res 43(3):e15.

49. Stamatakis A. 2014. RAxML version 8: A tool for phylogenetic analysis and post-analysis of large phylogenies. Bioinformatics. 30(9):1312–3.

50. Zhou Z, Alikhan NF, Sergeant MJ, Luhmann N, Vaz C, Francisco AP, Carriço AJ, Achtman M. 2018. Grapetree: Visualization of core genomic relationships among 100,000 bacterial pathogens. Genome Res 28(9):1395–404.

51. Altschul SF, Gish W, Miller W, Myers EW, Lipman DJ. 1990. Basic local alignment search tool. J Mol Biol 215(3):403–10.

52. Merda D, Felten A, Vingadassalon N, Denayer S, Titouche Y, Decastelli L, Hickey B, Kourtis C, Daskalov H, Mistou MY, Hennekinne JA. 2020. NAuRA: Genomic Tool to Identify Staphylococcal Enterotoxins in Staphylococcus aureus Strains Responsible for FoodBorne Outbreaks. Front Microbiol 11:1483

53. R Core Team. 2017. R: A Language and Environment for Statistical Computing. R Foundation for Statistical Computing, Vienna, Austria. https://www.R-project.org.

54. Bazzi AM, Rabaan AA, Fawarah MM, Al-Tawfiq JA. 2017. Direct identification and susceptibility testing of positive blood cultures using high speed cold centrifugation and Vitek II system. J Infect Public Health 10(3):299–307.

55. Holden MTG, Hsu LY, Kurt K, Weinert LA, Mather AE, Harris SR, Strommenger B, Layer F, Witte W, de Lencastre H, Skov R, Westh H, Zemlicková H, Coombs G, Kearns AM, Hill RLR, Edgeworth J, Gould I, Gant V, Cooke J, Edwards GF, McAdam PR, Zhou Z, Castillo-Ramírez S, Feil EJ, Hudson LO, Enright MC, Balloux F, Aanensen DM, Spratt BG, Fitzgerald RJ, Parkhill J, Achtman M, Bentley SD, Nübel U. 2013. A genomic portrait of the emergence, evolution, and global spread of a methicillin-resistant Staphylococcus aureus pandemic. Genome Res 23(4):653–64

56. Roe C, Stegger M, Lilje B, Johannesen TB, Ng KL, Sieber RN, Driebe E. Engelthaler DM, Andersen PS. 2020. Genomic analyses of Staphylococcus aureus clonal complex 45 isolates does not distinguish nasal carriage from bacteraemia. Microb Genom 6(8):1–9.

57. Saito E, Yoshida N, Kawano J, Shimizu A, Igimi S. 2011. Isolation of Staphylococcus aureus from raw fish in relation to culture methods. J Vet Med Sci 73(3):287–92.

58. British Trout Association. Trout Farming | British Trout Association. cited 2022 Aug 4. Available from: https://britishtrout.co.uk/about-trout/trout-farming/

59. Tolba O, Loughrey A, Goldsmith CE, Millar BC, Rooney PJ, Moore JE. 2008. Survival of epidemic strains of healthcare (HA-MRSA) and community-associated (CA-MRSA) meticillin-resistant Staphylococcus aureus (MRSA) in river-, sea- and swimming pool water. Int J Hyg Environ Health 211(3–4):398–402.

60. Levin-Edens E, Bonilla N, Meschke JS, Roberts MC. 2011. Survival of environmental and clinical strains of methicillin-resistant Staphylococcus aureus [MRSA] in marine and fresh waters. Water Res 45(17):5681–6.

61. EUCAST: Clinical breakpoints and dosing of antibiotics. cited 2022 Aug 4. Available from: https://www.eucast.org/clinical_breakpoints/

62. Cabello FC, Godfrey HP, Tomova A, Ivanova L, Dölz H, Millanao A, Buschmann AH. 2013. Antimicrobial use in aquaculture re-examined: its relevance to antimicrobial resistance and to animal and human health. Environ Microbiol 15(7):1917–42.

63. Use of Antimicrobial Agents in Aquaculture | The Fish Site. cited 2022 Aug 4. Available from: https://thefishsite.com/articles/use-of-antimicrobial-agents-in-aquaculture

64. van Wamel Wjb, Rooijakkers SHM, Ruyken M, van Kessel KPM, van Strijp Jag. 2006. The innate immune modulators staphylococcal complement inhibitor and chemotaxis inhibitory protein of Staphylococcus aureus are located on β-hemolysin-converting bacteriophages. J Bacteriol 188(4):1310–5.

65. Börjesson S, Matussek A, Melin S, Löfgren S, Lindgren PE. 2010. Methicillin-resistant Staphylococcus aureus (MRSA) in municipal wastewater: An uncharted threat? J Appl Microbiol 108(4):1244–1251.

66. Concepción Porrero M, Valverde A, Fernández-Llario P, Díez-Guerrier A, Mateos A, Lavín S, Cantón R, Fernández-Garayzabal JF, Domínguez L. 2014. Staphylococcus aureus carrying mecC gene in animals and Urban Wastewater, Emerg Infect Dis 20(5): 899–901.

67. Rosenberg Goldstein RE, Micallef SA, Gibbs SG, Davis JA, He X, George A, Kleinfelter LM, Schreiber NA, Mukherjee S, Sapkota A, Joseph SW, Sapkota AR. 2012. Methicillin-resistant Staphylococcus aureus (MRSA) Detected at Four U.S. wastewater treatment plants. Environ Health Perspect 120(11):1551–8.

68. Wan MT W, Chou CC. 2014. Spreading of β-lactam resistance gene (mecA) and methicillin-resistant Staphylococcus aureus through municipal and swine slaughterhouse wastewaters. Water Res 64:288–95.

69. Thompson JM, Gündoǧdu A, Stratton HM, Katouli M. 2013. Antibiotic resistant Staphylococcus aureus in hospital wastewaters and sewage treatment plants with special reference to methicillin-resistant Staphylococcus aureus (MRSA). J Appl Microbiol 114(1):44–54.

70. Brooks JP, Adeli A, McLaughlin MR. 2014. Microbial ecology, bacterial pathogens, and antibiotic resistant genes in swine manure wastewater as influenced by three swine management systems. Water Res 57:96–103.

71. Arfatahery N, Davoodabadi A, Abedimohtasab T. 2016. Characterization of Toxin Genes and Antimicrobial Susceptibility of Staphylococcus aureus Isolates in Fishery Products in Iran. Scientific Reports 6(1):1–7.

72. Rashid N, Shafee M, Iqbal S, Samad A, Khan SA, Hasni MS, Rehman ZU, Ullah S, Rehman FU, Khan GI, Ahmad S, Akbar A. 2021. Enterotoxigenic methicillin resistant Staphylococcus aureus contamination in salted fish from Gwadar Balochistan. Braz J Biol 83:e247701.

73. Hennekinne JA, de Buyser ML, Dragacci S. 2012. Staphylococcus aureus and its food poisoning toxins: characterization and outbreak investigation. FEMS Microbiol Rev 36(4):815–36.

74. Fisher EL, Otto M, Cheung GYC. 2018 Basis of virulence in enterotoxin-mediated staphylococcal food poisoning. Front Microbiol 9:436.

75. Bujjamma P, Padmavathi P. 2015. Prevalence of Staphylococcus aureus in Fish Samples of Local Domestic Fish Market. Int J Curr Microbiol Appl Sci 4(5):427–33.

76. Dong X, Wang X, Chen X, Yan Z, Cheng J, Gao L, Li Y, Li J. 2017. Genetic Diversity and Virulence Potential of Staphylococcus aureus Isolated from Crayfish (Procambarus clarkii). Curr Microbiol. 74(1):28–33.

77. Bank W. FISH TO 2030 Prospects for Fisheries and Aquaculture FISH TO 2030 Prospects for Fisheries and Aquaculture. 83177. Available from: https://openknowledge.worldbank.org/handle/10986/17579.

78. Blaylock RB, Bullard SA. 2014. Counter-insurgents of the blue revolution? parasites and diseases affecting aquaculture and science. J Parasitol 100(6):743–55.

79. Karvonen A, Rintamäki P, Jokela J, Valtonen ET. 2010. Increasing water temperature and disease risks in aquatic systems: Climate change increases the risk of some, but not all, diseases. Int J Parasitol 40(13):1483–8.

80. United Nations. 2020. Sustainable Development Goals Report – United Nations Sustainable Development. United Nations Publications. Available from: https://www.un.org/sustainabledevelopment/progress-report/

