## Supplemental Table 1 for "Absence of *Staphylococcus aureus* in wild populations of fish supports a spillover hypothesis"

**Table S1: Metadata for all CC45 *S. aureus* used in the phylogenetic reconstruction analysis.**

| *Strain* | *County* | *Host* | *ST* | *CC* | *ERR number* |
| --- | --- | --- | --- | --- | --- |
| 09.4504.Z | United Kingdom | Human | 45 | 45 | ERR084607 |
| 4238_1_1 | Belgium | Human | 45 | 45 | ERR033239 |
| 4238_1_10 | Belgium | Human | 45 | 45 | ERR033240 |
| 4238_1_7 | Belgium | Human | 45 | 45 | ERR033248 |
| 4238_1_8 | Belgium | Human | 45 | 45 | ERR033249 |
| 4330_8_1 | Austria | Human | 45 | 45 | ERR033304 |
| 4340_5_1 | Bulgaria | Human | 45 | 45 | ERR033330 |
| 4340_6_1 | Belgium | Human | 45 | 45 | ERR033343 |
| 4350_1_6 | Czech | Human | 45 | 45 | ERR033364 |
| 4386_1_11 | Croatia | Human | 45 | 45 | ERR033410 |
| 4386_1_2 | Croatia | Human | 45 | 45 | ERR033412 |
| 4386_1_3 | Cyprus | Human | 45 | 45 | ERR033413 |
| 4386_1_5 | Croatia | Human | 45 | 45 | ERR033415 |
| 4386_7_4 | Germany | Human | 45 | 45 | ERR033479 |
| 4395_3_11 | Spain | Human | 45 | 45 | ERR033526 |
| 4414_2_10 | Poland | Human | 45 | 45 | ERR033551 |
| 4414_2_6 | Poland | Human | 45 | 45 | ERR033558 |
| 4414_3_10 | Sweden | Human | 45 | 45 | ERR033564 |
| 4414_3_11 | Sweden | Human | 45 | 45 | ERR033565 |
| 4414_3_12 | Sweden | Human | 45 | 45 | ERR033566 |
| 4414_3_3 | Sweden | Human | 45 | 45 | ERR033568 |
| 4414_5_10 | Sweden | Human | 45 | 45 | ERR033577 |
| 4414_5_4 | Sweden | Human | 45 | 45 | ERR033582 |
| 4414_5_6 | Sweden | Human | 45 | 45 | ERR033584 |
| 4414_6_3 | Sweden | Human | 45 | 45 | ERR033594 |
| 4414_6_6 | Sweden | Human | 45 | 45 | ERR033597 |
| 4430_1_1 | Denmark | Human | 45 | 45 | ERR033628 |
| 4465_5_4 | Norway | Human | 45 | 45 | ERR033712 |
| 6133_3_5 | Poland | Human | 45 | 45 | ERR038686 |
| 6133_3_8 | Poland | Human | 45 | 45 | ERR038689 |
| 6236_1_4 | Austria | Human | 45 | 45 | ERR039391 |
| A-69 | Turkey | Cow | 45 | 45 | ERR387222 |
| B302368 | United Kingdom | Wild Bird | 45 | 45 | ERR2505672 |
| CSRJAE39 | Switzerland | Horse | 45 | 45 | ERR234859 |
| CTH14 | USA | Cow | 45 | 45 | ERR2505706 |
| CTH160 | USA | Cow | 45 | 45 | ERR2505707 |
| CTH211 | USA | Cow | 45 | 45 | ERR2505703 |
| Z_22 | Denmark | Horse | 45 | 45 | ERR2505688 |
| 4414_5_8 | Sweden | Human | 46 | 45 | ERR033586 |
| 4415_2_8 | United Kingdom | Human | 46 | 45 | ERR033612 |
| ASASM145 | United Kingdom | Human | 47 | 45 | ERR109591 |
| ASASM300 | United Kingdom | Human | 54 | 45 | ERR114905 |
| ASASM325 | United Kingdom | Human | 508 | 45 | ERR114924 |
| SA21 | Gambia | Human | 508 | 45 | ERR1213778 |
| 4386_2_1 | Denmark | Human | 682 | 45 | ERR033421 |
| 4350_1_7 | Czech | Human | 2860 | 45 | ERR033365 |
| 4386_2_6 | Denmark | Human | 2863 | 45 | ERR033429 |
| 6133_1_12 | France | Human | 2876 | 45 | ERR038669 |
| ASASM17 | United Kingdom | Human | 2902 | 45 | ERR109499 |
| ASASM158 | United Kingdom | Human | 2903 | 45 | ERR109571 |
| ASASM356 | United Kingdom | Human | 2904 | 45 | ERR109683 |
| ASASM171 | United Kingdom | Human | 2927 | 45 | ERR109616 |
| ASASM426 | United Kingdom | Human | 2940 | 45 | ERR172064 |
| 4330_8_4 | Belgium | Human | 3299 | 45 | ERR033310 |
| 45-164 | United Kingdom | Cow | 3613 | 45 | ERR294319 |
| 4350_1_4 | Czech | Human | 4670 | 45 | ERR033362 |
