## Supplemental Table 2 for "Absence of *Staphylococcus aureus* in wild populations of fish supports a spillover hypothesis"

**Table S2: MIC values obtained using Vitek2 for two rainbow trout isolates from fish farm in London.**

| *Antibiotic* | *BTGL0319001* | *BTIL0319003* |
| --- | --- | --- |
| Cefoxitin Screen | NEG | NEG |
| Benzylpenicillin | >=0.5 | >=0.5 |
| Amoxicillin/ Clavulanic acid | <=2 | <=2 |
| Oxacillin | <=0.25 | <=0.25 |
| Cefalotin | <=2 | <=2 |
| Cefovecin | 1 | 1 |
| Ceftiofur | 1 | 1 |
| Gentamicin | <=0.5 | <=0.5 |
| Kanamycin | <=4 | <=4 |
| Neomycin | <=2 | <=2 |
| Enrofloxacin | <=0.5 | <=0.5 |
| Marbofloxacin | <=0.5 | <=0.5 |
| Pradofloxacin | <=0.12 | <=0.12 |
| Inducible Clindamycin Resistance (ICR) | NEG | NEG |
| Erythromycin | <=0.25 | <=0.25 |
| Clindamycin | 0.25 | 0.25 |
| Doxycycline | <=0.5 | <=0.5 |
| Tetracycline | <=1 | <=1 |
| Nitrofurantoin | 32 | 32 |
| Chloramphenicol | 8 | <=4 |
| Trimethoprim / Sulfamethoxazole | <=10 | <=10 |
