## Supplemental Table 3 for "Absence of *Staphylococcus aureus* in wild populations of fish supports a spillover hypothesis"

**Table S4: Sampled lochs with sizes, coordinates.**

| *Sampling site* | *Location* | *Loch code* | *Perimeter (km)* | *Area (ha)* | *Altitude (m)* | *Habitat* | *Sampling date* |
| --- | --- | --- | --- | --- | --- | --- | --- |
| Loch a'Gharbh Doire | N57º44'52''  W5º41'51'' | LGD | 1.69 | 10.98 | 82 | Bird | 13 JUL 2019 |
| Loch Coire na h-Airigh | N57º44'29''  W5º41'25'' | LCA | 0.78 | 2.23 | 80 | Bird | 13 JUL 2019 |
| Loch Feur | N57º44'21''  W5º40'57'' | LFE | 1.16 | 4.55 | 75 | Bird | 13 JUL 2019 |
| Loch nam Breac | N57º44'27''  W5º40'30'' | LNB | 1.02 | 4.95 | 77 | Bird | 13 JUL 2019 |
| Greylag Loch | N57º54'40''  W5º33'36'' | GRB | 0.97 | 4.83 | 21 | Bird | 14 JUL 2019 |
| Loch Orchid | N57º37'47''  W5º16'55'' | GLO | 0.44 | 0.87 | 374 | Isolated | 12 JUL 2019 |
| Golden Loch A | N57º37'46''  W5º17'56'' | LFO | 0.63 | 1.61 | 420 | Isolated | 12 JUL 2019 |
| Golden Loch B | N57º37'44''  W5º17'29'' | LFI | 0.75 | 1 | 400 | Isolated | 12 JUL 2019 |
| Loch Dubh Dughaill | N57º42'18''  W5º36'02'' | LDD | 0.73 | 1.66 | 293 | Isolated | 15 JUL 2019 |
| Loch na Feithe Mugaig | N57º42'42''  W5º35'46'' | LFM | 3.65 | 16.27 | 306 | Isolated | 15 JUL 2019 |
| Loch nan Buainichean | N57º42'01''  W5º36'14'' | LB | 2.1 | 11.13 | 204 | Isolated | 15 JUL 2019 |
| Loch an Draing | N57º51'08''  W5º45'10'' | LDR | 3.16 | 38 | 41 | Livestock | 11 JUL 2019 |
| Loch nan Eun | N57º51'16''  W5º45'19'' | LEV | 2.15 | 16.46 | 40 | Livestock | 11 JUL 2019 |
| Loch na Fideil | N57º40'15''  W5º28'45'' | LNF | 0.62 | 2.5 | 21 | Livestock | 17 JUL 2019 |
| Loch Beag nan Eun | N57º51'30''  W5º45'39'' | LBE | 0.34 | 0.78 | 42 | Livestock | 19 JUL 2019 |
| Loch Bad an Scalaig | N57º41'11.12''W5º36'42.22'' | LBS |  |  | 114 | NA | 22 JUL 2019 |
| Allt Phadraig | N57º46'44''  W5º48'00'' | APD |  |  |  | NA | 19 JUL 2019 |
| Kerrysdale (River Kerry B8056 A832) | N57º41'35.34''W5º39'27.41'' | KED |  |  | 24 | NA | 22 JUL 2019 |
| Pollack Point | N57º54'45''  W5º33'05'' | SPP | NA | NA | NA | Sea | 14 JUL 2019 |
| Mellon Udrigle Bay | N57º54'12''  W5º33'22'' | MUB | NA | NA | NA | Sea | 14 JUL 2019 |
| River Canaird | N57º56'50''  W5º10'48'' | RCA | NA | NA | NA | Sea | 16 JUL 2019 |
| Sand River | N57º44'25''W5º46'19'' | SR | NA | NA | NA | Sea | 21 JUL 2019 |
| Flowerdale Estuary | N57º42'44''W5º40'43'' | FES | NA | NA | NA | Sea | 20 JUL 2019 |
| Sand Beach Ocean | N57º44'11''W5º46'07'' | SBO | NA | NA | NA | Sea | 21 JUL 2019 |
