## Supplemental Table 4 for "Absence of *Staphylococcus aureus* in wild populations of fish supports a spillover hypothesis"

**Table S5: Metadata for all samples collected from fish in Scottish Highlands. The loch code corresponds to names in the Table S3.**

| *Sample name* | *Tissue sampled* | *Loch code* | *Habitat* | *Host* | *Date of sampling* |
| --- | --- | --- | --- | --- | --- |
| BTG_LNB0719_001 | Gills | LNB | Bird Lochs | Brown Trout | 9 JUL 2019 |
| BTG_LNB0719_002 | Gills | LNB | Bird Lochs | Brown Trout | 9 JUL 2019 |
| BTG_LNB0719_003 | Gills | LNB | Bird Lochs | Brown Trout | 9 JUL 2019 |
| BTG_LNB0719_004 | Gills | LNB | Bird Lochs | Brown Trout | 9 JUL 2019 |
| BTG_LNB0719_005 | Gills | LNB | Bird Lochs | Brown Trout | 9 JUL 2019 |
| BTG_LNB0719_006 | Gills | LNB | Bird Lochs | Brown Trout | 9 JUL 2019 |
| BTG_LNB0719_007 | Gills | LNB | Bird Lochs | Brown Trout | 9 JUL 2019 |
| BTG_LNB0719_008 | Gills | LNB | Bird Lochs | Brown Trout | 9 JUL 2019 |
| BTG_LNB0719_009 | Gills | LNB | Bird Lochs | Brown Trout | 9 JUL 2019 |
| BTG_LNB0719_010 | Gills | LNB | Bird Lochs | Brown Trout | 9 JUL 2019 |
| BTV_LNB0719_001 | Vent | LNB | Bird Lochs | Brown Trout | 9 JUL 2019 |
| BTV_LNB0719_002 | Vent | LNB | Bird Lochs | Brown Trout | 9 JUL 2019 |
| BTV_LNB0719_003 | Vent | LNB | Bird Lochs | Brown Trout | 9 JUL 2019 |
| BTV_LNB0719_004 | Vent | LNB | Bird Lochs | Brown Trout | 9 JUL 2019 |
| BTV_LNB0719_005 | Vent | LNB | Bird Lochs | Brown Trout | 9 JUL 2019 |
| BTV_LNB0719_006 | Vent | LNB | Bird Lochs | Brown Trout | 9 JUL 2019 |
| BTV_LNB0719_007 | Vent | LNB | Bird Lochs | Brown Trout | 9 JUL 2019 |
| BTV_LNB0719_008 | Vent | LNB | Bird Lochs | Brown Trout | 9 JUL 2019 |
| BTV_LNB0719_009 | Vent | LNB | Bird Lochs | Brown Trout | 9 JUL 2019 |
| BTV_LNB0719_010 | Vent | LNB | Bird Lochs | Brown Trout | 9 JUL 2019 |
| BTG_LGD0719_001 | Gills | LGD | Bird Lochs | Brown Trout | 10 JUL 2019 |
| BTG_LGD0719_002 | Gills | LGD | Bird Lochs | Brown Trout | 10 JUL 2019 |
| BTG_LGD0719_003 | Gills | LGD | Bird Lochs | Brown Trout | 10 JUL 2019 |
| BTG_LGD0719_004 | Gills | LGD | Bird Lochs | Brown Trout | 10 JUL 2019 |
| BTG_LGD0719_005 | Gills | LGD | Bird Lochs | Brown Trout | 10 JUL 2019 |
| BTG_LGD0719_006 | Gills | LGD | Bird Lochs | Brown Trout | 10 JUL 2019 |
| BTG_LGD0719_007 | Gills | LGD | Bird Lochs | Brown Trout | 10 JUL 2019 |
| BTG_LGD0719_008 | Gills | LGD | Bird Lochs | Brown Trout | 10 JUL 2019 |
| BTG_LGD0719_009 | Gills | LGD | Bird Lochs | Brown Trout | 10 JUL 2019 |
| BTG_LGD0719_010 | Gills | LGD | Bird Lochs | Brown Trout | 10 JUL 2019 |
| BTG_LGD0719_011 | Gills | LGD | Bird Lochs | Brown Trout | 10 JUL 2019 |
| BTV_LGD0719_001 | Vent | LGD | Bird Lochs | Brown Trout | 10 JUL 2019 |
| BTV_LGD0719_002 | Vent | LGD | Bird Lochs | Brown Trout | 10 JUL 2019 |
| BTV_LGD0719_003 | Vent | LGD | Bird Lochs | Brown Trout | 10 JUL 2019 |
| BTV_LGD0719_004 | Vent | LGD | Bird Lochs | Brown Trout | 10 JUL 2019 |
| BTV_LGD0719_005 | Vent | LGD | Bird Lochs | Brown Trout | 10 JUL 2019 |
| BTV_LGD0719_006 | Vent | LGD | Bird Lochs | Brown Trout | 10 JUL 2019 |
| BTV_LGD0719_007 | Vent | LGD | Bird Lochs | Brown Trout | 10 JUL 2019 |
| BTV_LGD0719_008 | Vent | LGD | Bird Lochs | Brown Trout | 10 JUL 2019 |
| BTV_LGD0719_009 | Vent | LGD | Bird Lochs | Brown Trout | 10 JUL 2019 |
| BTV_LGD0719_010 | Vent | LGD | Bird Lochs | Brown Trout | 10 JUL 2019 |
| BTV_LGD0719_011 | Vent | LGD | Bird Lochs | Brown Trout | 10 JUL 2019 |
| BTG_LCA0719_001 | Gills | LCA | Bird Lochs | Brown Trout | 13 JUL 2019 |
| BTG_LCA0719_002 | Gills | LCA | Bird Lochs | Brown Trout | 13 JUL 2019 |
| BTG_LCA0719_003 | Gills | LCA | Bird Lochs | Brown Trout | 13 JUL 2019 |
| BTG_LCA0719_004 | Gills | LCA | Bird Lochs | Brown Trout | 13 JUL 2019 |
| BTG_LCA0719_005 | Gills | LCA | Bird Lochs | Brown Trout | 13 JUL 2019 |
| BTG_LCA0719_006 | Gills | LCA | Bird Lochs | Brown Trout | 13 JUL 2019 |
| BTG_LCA0719_007 | Gills | LCA | Bird Lochs | Brown Trout | 13 JUL 2019 |
| BTG_LCA0719_008 | Gills | LCA | Bird Lochs | Brown Trout | 13 JUL 2019 |
| BTG_LCA0719_009 | Gills | LCA | Bird Lochs | Brown Trout | 13 JUL 2019 |
| BTG_LCA0719_010 | Gills | LCA | Bird Lochs | Brown Trout | 13 JUL 2019 |
| BTG_LCA0719_011 | Gills | LCA | Bird Lochs | Brown Trout | 13 JUL 2019 |
| BTV_LCA0719_001 | Vent | LCA | Bird Lochs | Brown Trout | 13 JUL 2019 |
| BTV_LCA0719_002 | Vent | LCA | Bird Lochs | Brown Trout | 13 JUL 2019 |
| BTV_LCA0719_003 | Vent | LCA | Bird Lochs | Brown Trout | 13 JUL 2019 |
| BTV_LCA0719_004 | Vent | LCA | Bird Lochs | Brown Trout | 13 JUL 2019 |
| BTV_LCA0719_005 | Vent | LCA | Bird Lochs | Brown Trout | 13 JUL 2019 |
| BTV_LCA0719_006 | Vent | LCA | Bird Lochs | Brown Trout | 13 JUL 2019 |
| BTV_LCA0719_007 | Vent | LCA | Bird Lochs | Brown Trout | 13 JUL 2019 |
| BTV_LCA0719_008 | Vent | LCA | Bird Lochs | Brown Trout | 13 JUL 2019 |
| BTV_LCA0719_009 | Vent | LCA | Bird Lochs | Brown Trout | 13 JUL 2019 |
| BTV_LCA0719_010 | Vent | LCA | Bird Lochs | Brown Trout | 13 JUL 2019 |
| BTV_LCA0719_011 | Vent | LCA | Bird Lochs | Brown Trout | 13 JUL 2019 |
| BTG_LEF0719_001 | Gills | LFE | Bird Lochs | Brown Trout | 13 JUL 2019 |
| BTG_LEF0719_002 | Gills | LFE | Bird Lochs | Brown Trout | 13 JUL 2019 |
| BTG_LEF0719_003 | Gills | LFE | Bird Lochs | Brown Trout | 13 JUL 2019 |
| BTG_LEF0719_004 | Gills | LFE | Bird Lochs | Brown Trout | 13 JUL 2019 |
| BTG_LEF0719_005 | Gills | LFE | Bird Lochs | Brown Trout | 13 JUL 2019 |
| BTG_LEF0719_006 | Gills | LFE | Bird Lochs | Brown Trout | 13 JUL 2019 |
| BTG_LEF0719_007 | Gills | LFE | Bird Lochs | Brown Trout | 13 JUL 2019 |
| BTG_LEF0719_008 | Gills | LFE | Bird Lochs | Brown Trout | 13 JUL 2019 |
| BTG_LEF0719_009 | Gills | LFE | Bird Lochs | Brown Trout | 13 JUL 2019 |
| BTG_LEF0719_010 | Gills | LFE | Bird Lochs | Brown Trout | 13 JUL 2019 |
| BTV_LEF0719_001 | Vent | LFE | Bird Lochs | Brown Trout | 13 JUL 2019 |
| BTV_LEF0719_002 | Vent | LFE | Bird Lochs | Brown Trout | 13 JUL 2019 |
| BTV_LEF0719_003 | Vent | LFE | Bird Lochs | Brown Trout | 13 JUL 2019 |
| BTV_LEF0719_004 | Vent | LFE | Bird Lochs | Brown Trout | 13 JUL 2019 |
| BTV_LEF0719_005 | Vent | LFE | Bird Lochs | Brown Trout | 13 JUL 2019 |
| BTV_LEF0719_006 | Vent | LFE | Bird Lochs | Brown Trout | 13 JUL 2019 |
| BTV_LEF0719_007 | Vent | LFE | Bird Lochs | Brown Trout | 13 JUL 2019 |
| BTV_LEF0719_008 | Vent | LFE | Bird Lochs | Brown Trout | 13 JUL 2019 |
| BTV_LEF0719_009 | Vent | LFE | Bird Lochs | Brown Trout | 13 JUL 2019 |
| BTV_LEF0719_010 | Vent | LFE | Bird Lochs | Brown Trout | 13 JUL 2019 |
| BTG_LFO0719_001 | Gills | LFO | Isolated Lochs | Brown Trout | 12 JUL 2019 |
| BTG_LFO0719_002 | Gills | LFO | Isolated Lochs | Brown Trout | 12 JUL 2019 |
| BTG_LFO0719_003 | Gills | LFO | Isolated Lochs | Brown Trout | 12 JUL 2019 |
| BTG_LFO0719_004 | Gills | LFO | Isolated Lochs | Brown Trout | 12 JUL 2019 |
| BTG_LFO0719_005 | Gills | LFO | Isolated Lochs | Brown Trout | 12 JUL 2019 |
| BTG_LFO0719_006 | Gills | LFO | Isolated Lochs | Brown Trout | 12 JUL 2019 |
| BTG_LFO0719_007 | Gills | LFO | Isolated Lochs | Brown Trout | 12 JUL 2019 |
| BTG_LFI0719_001 | Gills | LFI | Isolated Lochs | Brown Trout | 12 JUL 2019 |
| BTG_LFI0719_002 | Gills | LFI | Isolated Lochs | Brown Trout | 12 JUL 2019 |
| BTG_GLO0719_001 | Gills | GLO | Isolated Lochs | Brown Trout | 12 JUL 2019 |
| BTG_GLO0719_002 | Gills | GLO | Isolated Lochs | Brown Trout | 12 JUL 2019 |
| BTG_GLO0719_003 | Gills | GLO | Isolated Lochs | Brown Trout | 12 JUL 2019 |
| BTG_GLO0719_004 | Gills | GLO | Isolated Lochs | Brown Trout | 12 JUL 2019 |
| BTG_GLO0719_005 | Gills | GLO | Isolated Lochs | Brown Trout | 12 JUL 2019 |
| BTG_GLO0719_006 | Gills | GLO | Isolated Lochs | Brown Trout | 12 JUL 2019 |
| BTG_GLO0719_007 | Gills | GLO | Isolated Lochs | Brown Trout | 12 JUL 2019 |
| BTG_GLO0719_008 | Gills | GLO | Isolated Lochs | Brown Trout | 12 JUL 2019 |
| BTG_GLO0719_009 | Gills | GLO | Isolated Lochs | Brown Trout | 12 JUL 2019 |
| BTG_GLO0719_010 | Gills | GLO | Isolated Lochs | Brown Trout | 12 JUL 2019 |
| BTG_LEM0719_001 | Gills | LFM | Isolated Lochs | Brown Trout | 15 JUL 2019 |
| BTG_LEM0719_002 | Gills | LFM | Isolated Lochs | Brown Trout | 15 JUL 2019 |
| BTG_LEM0719_003 | Gills | LFM | Isolated Lochs | Brown Trout | 15 JUL 2019 |
| BTG_LEM0719_004 | Gills | LFM | Isolated Lochs | Brown Trout | 15 JUL 2019 |
| BTG_LEM0719_005 | Gills | LFM | Isolated Lochs | Brown Trout | 15 JUL 2019 |
| BTG_LEM0719_006 | Gills | LFM | Isolated Lochs | Brown Trout | 15 JUL 2019 |
| BTG_LEM0719_007 | Gills | LFM | Isolated Lochs | Brown Trout | 15 JUL 2019 |
| BTG_LEM0719_008 | Gills | LFM | Isolated Lochs | Brown Trout | 15 JUL 2019 |
| BTG_LEM0719_009 | Gills | LFM | Isolated Lochs | Brown Trout | 15 JUL 2019 |
| BTG_LEM0719_010 | Gills | LFM | Isolated Lochs | Brown Trout | 15 JUL 2019 |
| BTG_LEM0719_011 | Gills | LFM | Isolated Lochs | Brown Trout | 15 JUL 2019 |
| BTG_LEM0719_012 | Gills | LFM | Isolated Lochs | Brown Trout | 15 JUL 2019 |
| BTG_LEM0719_013 | Gills | LFM | Isolated Lochs | Brown Trout | 15 JUL 2019 |
| BTG_LEM0719_014 | Gills | LFM | Isolated Lochs | Brown Trout | 15 JUL 2019 |
| BTG_LEM0719_015 | Gills | LFM | Isolated Lochs | Brown Trout | 15 JUL 2019 |
| BTG_LEM0719_016 | Gills | LFM | Isolated Lochs | Brown Trout | 15 JUL 2019 |
| BTG_LEM0719_017 | Gills | LFM | Isolated Lochs | Brown Trout | 15 JUL 2019 |
| BTG_LEM0719_018 | Gills | LFM | Isolated Lochs | Brown Trout | 15 JUL 2019 |
| BTG_LEM0719_019 | Gills | LFM | Isolated Lochs | Brown Trout | 15 JUL 2019 |
| BTG_LEM0719_020 | Gills | LFM | Isolated Lochs | Brown Trout | 15 JUL 2019 |
| BTG_LEM0719_021 | Gills | LFM | Isolated Lochs | Brown Trout | 15 JUL 2019 |
| BTV_LEM0719_001 | Gills | LFM | Isolated Lochs | Brown Trout | 15 JUL 2019 |
| BTG_LDD0719_001 | Gills | LDD | Isolated Lochs | Brown Trout | 15 JUL 2019 |
| BTG_LDD0719_002 | Gills | LDD | Isolated Lochs | Brown Trout | 15 JUL 2019 |
| BTG_LDD0719_003 | Gills | LDD | Isolated Lochs | Brown Trout | 15 JUL 2019 |
| BTG_LDD0719_004 | Gills | LDD | Isolated Lochs | Brown Trout | 15 JUL 2019 |
| BTG_LDD0719_005 | Gills | LDD | Isolated Lochs | Brown Trout | 15 JUL 2019 |
| BTG_LDD0719_006 | Gills | LDD | Isolated Lochs | Brown Trout | 15 JUL 2019 |
| BTG_LDD0719_007 | Gills | LDD | Isolated Lochs | Brown Trout | 15 JUL 2019 |
| BTG_LDD0719_008 | Gills | LDD | Isolated Lochs | Brown Trout | 15 JUL 2019 |
| BTV_LFO0719_001 | Vent | LFO | Isolated Lochs | Brown Trout | 12 JUL 2019 |
| BTV_LFO0719_002 | Vent | LFO | Isolated Lochs | Brown Trout | 12 JUL 2019 |
| BTV_LFO0719_003 | Vent | LFO | Isolated Lochs | Brown Trout | 12 JUL 2019 |
| BTV_LFO0719_004 | Vent | LFO | Isolated Lochs | Brown Trout | 12 JUL 2019 |
| BTV_LFO0719_005 | Vent | LFO | Isolated Lochs | Brown Trout | 12 JUL 2019 |
| BTV_LFO0719_006 | Vent | LFO | Isolated Lochs | Brown Trout | 12 JUL 2019 |
| BTV_LFO0719_007 | Vent | LFO | Isolated Lochs | Brown Trout | 12 JUL 2019 |
| BTV_LFI0719_001 | Vent | LFI | Isolated Lochs | Brown Trout | 12 JUL 2019 |
| BTV_LFI0719_002 | Vent | LFI | Isolated Lochs | Brown Trout | 12 JUL 2019 |
| BTV_GLO0719_001 | Vent | GLO | Isolated Lochs | Brown Trout | 12 JUL 2019 |
| BTV_GLO0719_002 | Vent | GLO | Isolated Lochs | Brown Trout | 12 JUL 2019 |
| BTV_GLO0719_003 | Vent | GLO | Isolated Lochs | Brown Trout | 12 JUL 2019 |
| BTV_GLO0719_004 | Vent | GLO | Isolated Lochs | Brown Trout | 12 JUL 2019 |
| BTV_GLO0719_005 | Vent | GLO | Isolated Lochs | Brown Trout | 12 JUL 2019 |
| BTV_GLO0719_006 | Vent | GLO | Isolated Lochs | Brown Trout | 12 JUL 2019 |
| BTV_GLO0719_007 | Vent | GLO | Isolated Lochs | Brown Trout | 12 JUL 2019 |
| BTV_GLO0719_008 | Vent | GLO | Isolated Lochs | Brown Trout | 12 JUL 2019 |
| BTV_GLO0719_009 | Vent | GLO | Isolated Lochs | Brown Trout | 12 JUL 2019 |
| BTV_GLO0719_010 | Vent | GLO | Isolated Lochs | Brown Trout | 12 JUL 2019 |
| BTV_LEM0719_002 | Vent | LFM | Isolated Lochs | Brown Trout | 15 JUL 2019 |
| BTV_LEM0719_003 | Vent | LFM | Isolated Lochs | Brown Trout | 15 JUL 2019 |
| BTV_LEM0719_004 | Vent | LFM | Isolated Lochs | Brown Trout | 15 JUL 2019 |
| BTV_LEM0719_005 | Vent | LFM | Isolated Lochs | Brown Trout | 15 JUL 2019 |
| BTV_LEM0719_006 | Vent | LFM | Isolated Lochs | Brown Trout | 15 JUL 2019 |
| BTV_LEM0719_007 | Vent | LFM | Isolated Lochs | Brown Trout | 15 JUL 2019 |
| BTV_LEM0719_008 | Vent | LFM | Isolated Lochs | Brown Trout | 15 JUL 2019 |
| BTV_LEM0719_009 | Vent | LFM | Isolated Lochs | Brown Trout | 15 JUL 2019 |
| BTV_LEM0719_010 | Vent | LFM | Isolated Lochs | Brown Trout | 15 JUL 2019 |
| BTV_LEM0719_011 | Vent | LFM | Isolated Lochs | Brown Trout | 15 JUL 2019 |
| BTV_LEM0719_012 | Vent | LFM | Isolated Lochs | Brown Trout | 15 JUL 2019 |
| BTV_LEM0719_013 | Vent | LFM | Isolated Lochs | Brown Trout | 15 JUL 2019 |
| BTV_LEM0719_014 | Vent | LFM | Isolated Lochs | Brown Trout | 15 JUL 2019 |
| BTV_LEM0719_015 | Vent | LFM | Isolated Lochs | Brown Trout | 15 JUL 2019 |
| BTV_LEM0719_016 | Vent | LFM | Isolated Lochs | Brown Trout | 15 JUL 2019 |
| BTV_LEM0719_017 | Vent | LFM | Isolated Lochs | Brown Trout | 15 JUL 2019 |
| BTV_LEM0719_018 | Vent | LFM | Isolated Lochs | Brown Trout | 15 JUL 2019 |
| BTV_LEM0719_019 | Vent | LFM | Isolated Lochs | Brown Trout | 15 JUL 2019 |
| BTV_LEM0719_020 | Vent | LFM | Isolated Lochs | Brown Trout | 15 JUL 2019 |
| BTV_LEM0719_021 | Vent | LFM | Isolated Lochs | Brown Trout | 15 JUL 2019 |
| BTV_LDD0719_001 | Vent | LDD | Isolated Lochs | Brown Trout | 15 JUL 2019 |
| BTV_LDD0719_002 | Vent | LDD | Isolated Lochs | Brown Trout | 15 JUL 2019 |
| BTV_LDD0719_003 | Vent | LDD | Isolated Lochs | Brown Trout | 15 JUL 2019 |
| BTV_LDD0719_004 | Vent | LDD | Isolated Lochs | Brown Trout | 15 JUL 2019 |
| BTV_LDD0719_005 | Vent | LDD | Isolated Lochs | Brown Trout | 15 JUL 2019 |
| BTV_LDD0719_006 | Vent | LDD | Isolated Lochs | Brown Trout | 15 JUL 2019 |
| BTV_LDD0719_007 | Vent | LDD | Isolated Lochs | Brown Trout | 15 JUL 2019 |
| BTV_LDD0719_008 | Vent | LDD | Isolated Lochs | Brown Trout | 15 JUL 2019 |
| BTG_LDR0719_001 | Gills | LDR | Livestock Lochs | Brown Trout | 11 JUL 2019 |
| BTG_LDR0719_002 | Gills | LDR | Livestock Lochs | Brown Trout | 11 JUL 2019 |
| BTV_LDR0719_001 | Vent | LDR | Livestock Lochs | Brown Trout | 11 JUL 2019 |
| BTV_LDR0719_002 | Vent | LDR | Livestock Lochs | Brown Trout | 11 JUL 2019 |
| BTG_LBE0719_001 | Gills | LBE | Livestock Lochs | Brown Trout | 19 JUN 2019 |
| BTV_LBE0719_001 | Vent | LBE | Livestock Lochs | Brown Trout | 19 JUN 2019 |
| BTG_LNF0719_001 | Gills | LNF | Livestock Lochs | Brown Trout | 17 JUL 2019 |
| BTG_LNF0719_002 | Gills | LNF | Livestock Lochs | Brown Trout | 17 JUL 2019 |
| BTG_LNF0719_003 | Gills | LNF | Livestock Lochs | Brown Trout | 17 JUL 2019 |
| BTG_LNF0719_004 | Gills | LNF | Livestock Lochs | Brown Trout | 17 JUL 2019 |
| BTG_LNF0719_005 | Gills | LNF | Livestock Lochs | Brown Trout | 17 JUL 2019 |
| BTG_LNF0719_006 | Gills | LNF | Livestock Lochs | Brown Trout | 17 JUL 2019 |
| BTG_LNF0719_007 | Gills | LNF | Livestock Lochs | Brown Trout | 17 JUL 2019 |
| BTG_LNF0719_008 | Gills | LNF | Livestock Lochs | Brown Trout | 17 JUL 2019 |
| BTG_LNF0719_009 | Gills | LNF | Livestock Lochs | Brown Trout | 17 JUL 2019 |
| BTV_LNF0719_001 | Vent | LNF | Livestock Lochs | Brown Trout | 17 JUL 2019 |
| BTV_LNF0719_002 | Vent | LNF | Livestock Lochs | Brown Trout | 17 JUL 2019 |
| BTV_LNF0719_003 | Vent | LNF | Livestock Lochs | Brown Trout | 17 JUL 2019 |
| BTV_LNF0719_004 | Vent | LNF | Livestock Lochs | Brown Trout | 17 JUL 2019 |
| BTV_LNF0719_005 | Vent | LNF | Livestock Lochs | Brown Trout | 17 JUL 2019 |
| BTV_LNF0719_006 | Vent | LNF | Livestock Lochs | Brown Trout | 17 JUL 2019 |
| BTV_LNF0719_007 | Vent | LNF | Livestock Lochs | Brown Trout | 17 JUL 2019 |
| BTV_LNF0719_008 | Vent | LNF | Livestock Lochs | Brown Trout | 17 JUL 2019 |
| BTV_LNF0719_009 | Vent | LNF | Livestock Lochs | Brown Trout | 17 JUL 2019 |
| BTG_LEV0719_001 | Gills | LEU | Livestock Lochs | Brown Trout | 11 JUL 2019 |
| BTG_LEV0719_002 | Gills | LEU | Livestock Lochs | Brown Trout | 11 JUL 2019 |
| BTG_LEV0719_003 | Gills | LEU | Livestock Lochs | Brown Trout | 11 JUL 2019 |
| BTG_LEV0719_004 | Gills | LEV | Livestock Lochs | Brown Trout | 19 JUL 2019 |
| BTG_LEV0719_005 | Gills | LEV | Livestock Lochs | Brown Trout | 19 JUL 2019 |
| BTG_LEV0719_006 | Gills | LEV | Livestock Lochs | Brown Trout | 19 JUL 2019 |
| BTG_LEV0719_007 | Gills | LEV | Livestock Lochs | Brown Trout | 19 JUL 2019 |
| BTG_LEV0719_008 | Gills | LEV | Livestock Lochs | Brown Trout | 19 JUL 2019 |
| BTG_LEV0719_009 | Gills | LEV | Livestock Lochs | Brown Trout | 19 JUL 2019 |
| BTG_LEV0719_010 | Gills | LEV | Livestock Lochs | Brown Trout | 19 JUL 2019 |
| BTG_LEV0719_011 | Gills | LEV | Livestock Lochs | Brown Trout | 19 JUL 2019 |
| BTG_LEV0719_012 | Gills | LEV | Livestock Lochs | Brown Trout | 19 JUL 2019 |
| BTG_LEV0719_013 | Gills | LEV | Livestock Lochs | Brown Trout | 19 JUL 2019 |
| BTG_LEV0719_014 | Gills | LEV | Livestock Lochs | Brown Trout | 19 JUL 2019 |
| BTG_LEV0719_015 | Gills | LEV | Livestock Lochs | Brown Trout | 19 JUL 2019 |
| BTG_LEV0719_016 | Gills | LEV | Livestock Lochs | Brown Trout | 19 JUL 2019 |
| BTG_LEV0719_017 | Gills | LEV | Livestock Lochs | Brown Trout | 19 JUL 2019 |
| BTG_LEV0719_018 | Gills | LEV | Livestock Lochs | Brown Trout | 19 JUL 2019 |
| BTV_LEV0719_001 | Vent | LEU | Livestock Lochs | Brown Trout | 11 JUL 2019 |
| BTV_LEV0719_002 | Vent | LEU | Livestock Lochs | Brown Trout | 11 JUL 2019 |
| BTV_LEV0719_003 | Vent | LEU | Livestock Lochs | Brown Trout | 11 JUL 2019 |
| BTV_LEV0719_004 | Vent | LEV | Livestock Lochs | Brown Trout | 19 JUL 2019 |
| BTV_LEV0719_005 | Vent | LEV | Livestock Lochs | Brown Trout | 19 JUL 2019 |
| BTV_LEV0719_006 | Vent | LEV | Livestock Lochs | Brown Trout | 19 JUL 2019 |
| BTV_LEV0719_007 | Vent | LEV | Livestock Lochs | Brown Trout | 19 JUL 2019 |
| BTV_LEV0719_008 | Vent | LEV | Livestock Lochs | Brown Trout | 19 JUL 2019 |
| BTV_LEV0719_009 | Vent | LEV | Livestock Lochs | Brown Trout | 19 JUL 2019 |
| BTV_LEV0719_010 | Vent | LEV | Livestock Lochs | Brown Trout | 19 JUL 2019 |
| BTV_LEV0719_011 | Vent | LEV | Livestock Lochs | Brown Trout | 19 JUL 2019 |
| BTV_LEV0719_012 | Vent | LEV | Livestock Lochs | Brown Trout | 19 JUL 2019 |
| BTV_LEV0719_013 | Vent | LEV | Livestock Lochs | Brown Trout | 19 JUL 2019 |
| BTV_LEV0719_014 | Vent | LEV | Livestock Lochs | Brown Trout | 19 JUL 2019 |
| BTV_LEV0719_015 | Vent | LEV | Livestock Lochs | Brown Trout | 19 JUL 2019 |
| BTV_LEV0719_016 | Vent | LEV | Livestock Lochs | Brown Trout | 19 JUL 2019 |
| BTV_LEV0719_017 | Vent | LEV | Livestock Lochs | Brown Trout | 19 JUL 2019 |
| BTV_LEV0719_018 | Vent | LEV | Livestock Lochs | Brown Trout | 19 JUL 2019 |
| BTG_RSA0719_001 | Gills | RCA | Sea | Brown Trout | 16 JUL 2019 |
| BTG_RSA0719_002 | Gills | RCA | Sea | Brown Trout | 16 JUL 2019 |
| BTG_SRI0719_001 | Gills | SR | Sea | Brown Trout | 18 JUL 2019 |
| BTV_RSA0719_001 | Vent | RCA | Sea | Brown Trout | 16 JUL 2019 |
| BTV_RSA0719_002 | Vent | RCA | Sea | Brown Trout | 16 JUL 2019 |
| BTV_SRI0719_001 | Vent | SR | Sea | Brown Trout | 18 JUL 2019 |
