## Supplemental Table 5 for "Absence of *Staphylococcus aureus* in wild populations of fish supports a spillover hypothesis"

**Table S6: Metadata for all environmental samples collected in Scottish Highlands. The loch code corresponds to names in the Table S3.**

| *Loch code* | *Habitat* | *Date of sampling* |
| --- | --- | --- |
| LGD | Bird | 13_JUL_2019 |
| LCA | Bird | 13_JUL_2019 |
| LFE | Bird | 13_JUL_2019 |
| LNB | Bird | 13_JUL_2019 |
| GRB | Bird | 14_JUL_2019 |
| GLO | Isolated | 12_JUL_2019 |
| LFO | Isolated | 12_JUL_2019 |
| LFI | Isolated | 12_JUL_2019 |
| LMG | Isolated | 12_JUL_2019 |
| LDD | Isolated | 15_JUL_2019 |
| LFM | Isolated | 15_JUL_2019 |
| LB | Isolated | 15_JUL_2019 |
| LDR | Livestock | 11_JUL_2019 |
| LEU | Livestock | 11_JUL_2019 |
| LNF | Livestock | 17_JUL_2019 |
| LBE | Livestock | 19_JUL_2019 |
| LBS | Loch Additional sample | 22_JUL_2019 |
| APD | River Additional sample | 19_JUL_2019 |
| KED | River Additional sample | 22_JUL_2019 |
| SPP | Sea | 14_JUL_2019 |
| MUB | Sea | 14_JUL_2019 |
| RCA | Sea | 16_JUL_2019 |
| SR | Sea | 21_JUL_2019 |
| FES | Sea | 20_JUL_2019 |
| SBO | Sea | 21_JUL_2019 |
