## Supplemental Table 6 for "Absence of *Staphylococcus aureus* in wild populations of fish supports a spillover hypothesis"

**Table S7 Metadata and Sample Accession for all the isolates used in this study.**

| *Strain* | *Sampling site* | *Sampling type* | *ST* | *CC* | *Sample Accession* | *ERR number* |
| --- | --- | --- | --- | --- | --- | --- |
| BTIL0319_003 | Intestine | Swabbing | 54 | 45 | SAMEA5419346 | ERR4188682 |
| BLSL0319_003 | Skin | Swabbing | 54 | 45 | SAMEA5419347 | ERR4188685 |
| BLGL0319_003 | Gill | Tissue | 54 | 45 | SAMEA5419348 | ERR4188688 |
| BTIL0319_001 | Intestine | Tissue | 54 | 45 | SAMEA5419349 | ERR4188691 |
| BLGL0319_002 | Gill | Swabbing | 54 | 45 | SAMEA5419350 | ERR4188694 |
| BLSL0319_002 | Skin | Swabbing | 54 | 45 | SAMEA5419351 | ERR4188697 |
| BLGL0319_004 | Gill | Tissue | 54 | 45 | SAMEA5419352 | ERR4188700 |
| BLSL0319_004 | Skin | Tissue | 54 | 45 | SAMEA5419353 | ERR4188703 |
| BTIL0319_004 | Intestine | Tissue | 54 | 45 | SAMEA5419354 | ERR4188705 |
| BTIL0319_002 | Intestine | Swabbing | 54 | 45 | SAMEA5419355 | ERR4188708 |
| BLGL0319_001 | Gill | Swabbing | 54 | 45 | SAMEA5419356 | ERR4188711 |
| BLSL0319_001 | Skin | Tissue | 54 | 45 | SAMEA5419357 | ERR4188714 |
